# Fibroblast Nrf2 inhibits profibrotic transcription with Ddx54 and mitigates pathological fibrosis in the mouse heart and kidney

**DOI:** 10.1101/2023.11.09.566496

**Authors:** Toshiyuki Nishiji, Atsushi Hoshino, Nariko IKemura, Mellissa C Alcantara, Yusuke Higuchi, Yuki Uchio, Yuhei Kirita, Shunta Taminishi, Yuki Ozaki, Daisuke Motooka, Ryosuke Kobayashi, Takuro Horii, Izuho Hatada, Satoaki Matoba

## Abstract

**Background:** Tissue fibrosis is a common feature of many organ dysfunctions, such as heart failure and chronic kidney disease. However, no fundamental treatment has been developed. This study aims to identify novel molecular mechanisms for antifibrotic intervention, focusing on fibroblast activation.

**Methods:** We performed a forward genetic screen using a genome-wide CRISPR library in the context of transforming growth factor β (TGF-β)-mediated connective tissue growth factor (CTGF) expression, and used unbiased techniques such as Cleavage Under Targets and Tagmentation (CUT&Tag) and proximity-dependent biotin labeling by TurboID to reveal the detailed molecular mechanisms.

**Results:** CRISPR library screening identified a number of players in both the canonical Smad pathway and the non-canonical pathway. In addition to the known factors, the Keap1-Nrf2 pathway was identified as a predominant regulator of TGF-β-mediated CTGF expression. *Keap1* deletion and consequent Nrf2 activation broadly suppressed profibrotic gene expression, independently of conventional antioxidant effects. CUT&Tag revealed that Nrf2 bound to the proximity of fibrosis-related genes including *Ctgf* and *Fn1.* Subsequent individual analysis revealed Smad3 and RNA polymerase II binding to the Nrf2 peak site, which was attenuated by *Keap1* deletion. TurboID experiments further discovered that Nrf2 interacts with Ddx54, which acts as a corepressor. Consistently, *Keap1* deletion-mediated repression of profibrotic gene expression was reversed by additional *Ddx54* deletion. The impact of the Keap1-Nrf2 pathway on pathological fibrosis was examined using tamoxifen-inducible fibroblast-specific *Keap1* knockout mice. Pressure overload for 4 weeks robustly induced cardiac hypertrophy, fibrosis and contractile dysfunction. However, deletion of *Keap1* in the *Postn* lineage attenuated these cardiac pathologies. The anti-fibrotic effects of Keap1 deletion were also confirmed in renal fibrosis in the unilateral ureteral obstruction (UUO) model.

**Conclusions:** Fibroblast Nrf2 transcriptionally represses fibrosis-related genes in cooperation with the corepressor Ddx54. Fibroblast-specific deletion of *Keap1* attenuated pathological fibrosis in pressure overload heart failure and renal fibrosis.

## Introduction

Fibrosis is a pathological process characterized by the excessive accumulation of extracellular matrix (ECM) components such as collagen and fibronectin in tissues^1^. It is a common feature of various pathologies such as heart failure, chronic kidney disease, pulmonary fibrosis, liver cirrhosis and tumors, and is associated with up to 45% of all deaths in the developed world^2^. Fibroblasts are the most abundant cell type of the connective tissues found throughout the body and become activated and then differentiated into myofibroblasts^3^. Upon tissue injury, local tissue fibroblasts are activated, secreting inflammatory mediators and synthesizing ECM components. Myofibroblasts are also equipped with a contractile apparatus through the expression of α-smooth muscle actin (αSMA), which allows them to physically manipulate ECM fibers to close open wounds^4^. When tissue damage is minimal and non-recurrent, wound healing is efficient and deposition of ECM components is transient, resulting in the restoration of functional tissue architecture. However, with repetitive or severe injury, accumulation of ECM components continues, resulting in disruption of tissue architecture and organ dysfunction^1^.

Transforming growth factor β (TGF-β) is the primary driver of fibrosis in injured or diseased tissue. It acts on fibroblasts and myofibroblasts to promote proliferation, migration, matrix production, production of chemotactic signals that promote leukocyte recruitment, fibrosis, and differentiation of fibroblasts to myofibroblasts^3^. TGF-β signals through a heterodimeric receptor composed of two types of transmembrane serine/threonine kinase receptors, type I (TβRI) and type II (TβRII). The canonical pathway downstream of TGF-β involves phosphorylation of Smad2 and Smad3 by the heterodimeric receptor. The phosphorylated Smad2/3 then form a complex with Smad4 and translocate to the nucleus where they regulate gene expression. The non-canonical pathway involves the activation of other signaling pathways such as MAPK (ERK, p38, JNK), PI3K/Akt, Rho GTPase pathways^5–7^. TGF-β is a multifunctional cytokine. In addition to fibrosis, it regulates cell growth, differentiation, and apoptosis in various tissues and cells, including the immune system. Therefore, direct intervention on TGF-β is unlikely to be therapeutically feasible and other targets are required to control pathological fibrosis^3^.

To further understand the molecular mechanism of the profibrotic response to TGF-β signaling, in this study we performed a genome-wide CRISPR library screen and identified Keap1-Nrf2 as a key regulatory pathway. Essentially, the Keap1-Nrf2 pathway plays a critical role in cellular defense against xenobiotic and oxidative stress. Keap1 functions as a substrate adaptor protein for cullin 3 E3 ligase complexes and binds to Nrf2 for the ubiquitin proteasome system. When the Keap1 protein senses oxidative stress or electrophilic compounds, structural changes due to covalent bonding cause the release of the transcription factor Nrf2 from its inhibitory complex. Nrf2 then translocates to the nucleus and binds to antioxidant response elements (AREs) in the promoter regions of target genes, such as antioxidant and detoxification genes^8^. Recent evidence suggests broader regulation including inhibition of proinflammatory cytokine transcription^9,10^. This study extended the regulatory scope of the Keap1-NRF2 system to include profibrotic gene suppression upon TGF-β treatment, which was achieved in cooperation with the transcriptional corepressor Ddx54. When Keap1 was deleted in myofibroblasts, fibrotic pathology was attenuated in the pressure overload heart failure and chronic kidney disease, suggesting that intervention in the Keap1-Nrf2 system holds promise for the treatment of fibrotic tissue dysfunction.

## Results

### Genome-wide CRISPR-Cas9 knockout screen to dissect regulatory networks of the TGF-β-mediated profibrotic response

To understand the detailed mechanism between TGF-β and fibrosis, we performed a genome-wide CRISPR-Cas9 knockout screen. When treated with TGF-β, mouse mesenchymal progenitor C2C12 cells expressed representative genes associated with fibrosis. Among them, connective tissue growth factor (CTGF) showed a greater response to TGF-β than others, we decided to use CTGF expression as a reporter of profibrotic activation (Figure S1A). To develop a screening assay, we generated *Ctgf* knock-in reporter cells. The *P2A-GFP-loxP-pPGK-puro-loxP* fragment was inserted just before the stop codon of the *Ctgf* gene using the CRISPR-Cas9 system, and the *pPGK-puro* cassette was removed by adenovirus-mediated Cre expression (Figure 1A). We obtained 5 clones of *Ctgf-P2A-GFP* reporter and selected the clone 23 showing better and response to TGF-β treatment, which was similar with ∼15-fold upregulation in endogenous CTGF expression (Figure S1B). We first introduced Cas9 into these reporter cells, and the resulting Cas9-expressing *Ctgf-P2A-GFP* cells were infected with validated lentiviral particles generated from a whole-genome CRISPR-Brie library^11^. The pooled reporter cells expressing the library of gene-specific gRNAs were then subjected to TGF-β treatment for 24 hours to activate the profibrotic response. The highest and lowest 20% of cells with GFP signal were collected by fluorescence activated cell sorting (FACS) for genomic DNA isolation and further analysis (Figure 1B). We systematically analyzed genes whose gRNAs affected CTGF expression using the MAGeCK algorithm. In addition to *Ctgf*, several known components of the canonical TGF-β pathway, including *Tgfbr1,2* and *Smad2,3,4,7* were identified as high-confidence hits in the *Ctgf-P2A-GFP* reporter cells, supporting the robustness of this screen (Figure 1C). Gene Set Enrichment Analysis (GSEA) of gRNAs attenuating CTGF expression showed enrichment of gene sets associated with the TGF-β-Smad pathway and non-canonical MAPK and JNK pathways, as well as autophagy, which can be activated by TGF-β and promotes collagen release and consequent profibrotic response^12^. Meanwhile, several gene sets associated with epigenetic regulations were identified for CTGF upregulation (Figure 1D).

**Figure 1.**
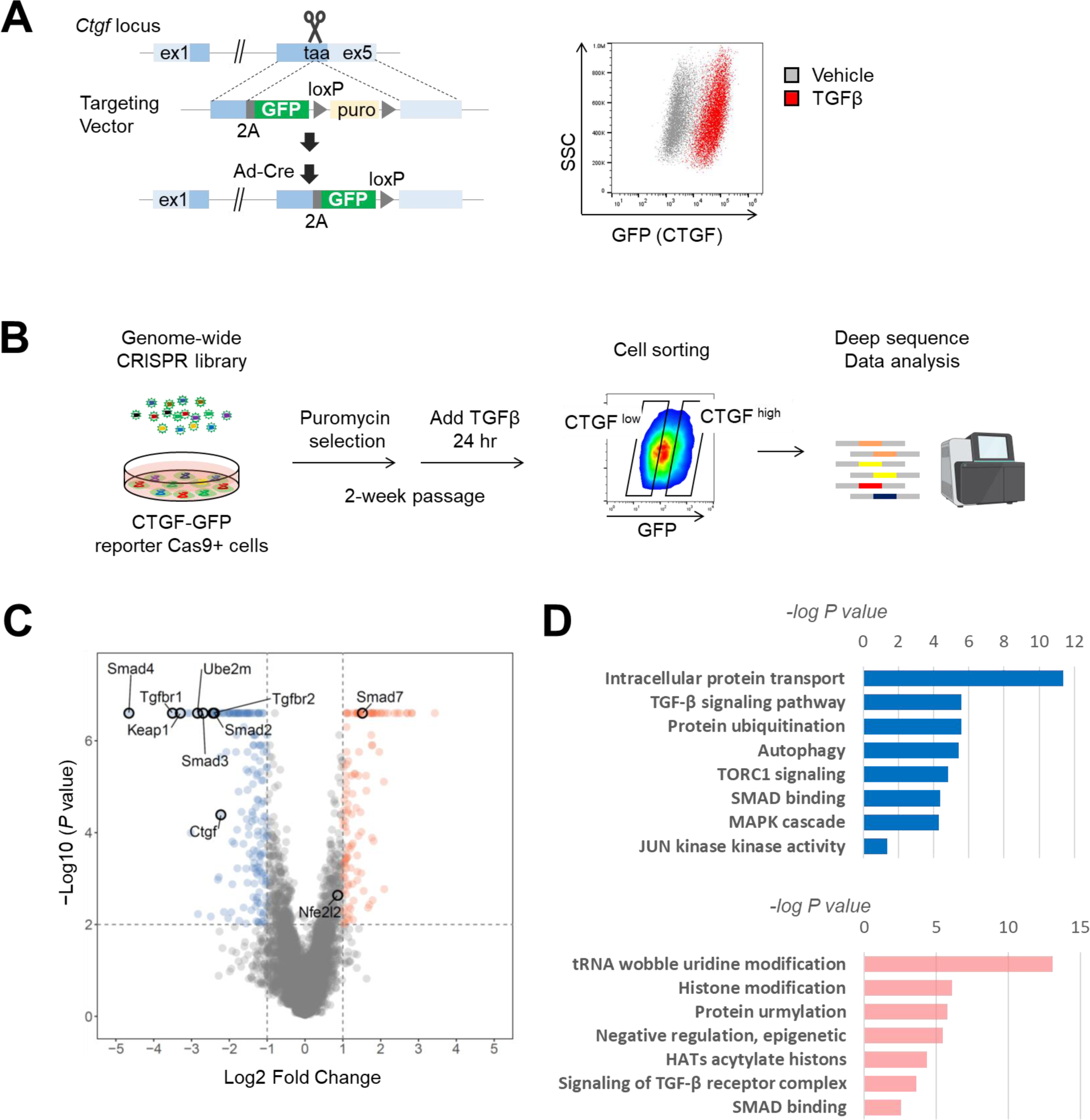
Genome-wide CRISPR-Cas9 knockout screen identifies genes that regulate TGF-β-mediated CTGF expression. **A**, The CTGF knock-in reporter cells were generated with mouse C2C12 mesenchymal progenitor cells using the CRISPR-Cas9 system. The *P2A-GFP-loxP-pPGK-puro-loxP* fragment was inserted just before the stop codon of *Ctgf* gene, and the *pPGK-puro* cassette was removed by adenovirus-mediated Cre expression (**left**). The resultant reporter cells were confirmed to express GFP upon treatment with TGF-β (10 ng/mL) (**right**). **B**, Schematic of genome-wide CRISPR screening using CTGF reporter cells. **C**, Systematic analysis of genes whose gRNAs affect CTGF expression based on the MAGeCK algorithm. **D**, Results of Gene Set Enrichment Analysis (GSEA) for gRNAs that substantially decrease (**top**) or increase (**bottom**) CTGF expression.

Focusing on the TGF-β pathway in the Kyoto Encyclopedia of Genes and Genomes (KEGG) pathway database (https://www.genome.jp/kegg/), TβRII activates TβRI to bind to Smad2 and Smad3, which then become active and associate with Smad4. Smad7 is the inhibitory Smad that can interact with activated TβRI and recruit the HECT domain-containing E3 ligases Smurf1 and Smurf2 to the receptor, leading to ubiquitination and degradation^13^. These factors were all identified (Figure 2A). In the step of Smad complex translocation into the nucleus, this study identified Xpo6, Xpo7, and Ipo8 as regulators, although Ipo7, Ipo8, and Xpo4 were previously reported to be associated with nucleocytoplasmic trafficking of Smad^14^. Among the non-canonical signaling pathways, the MAPK pathway intricately regulates fibrosis upon TGF-β, where genes in the JNK1/2 pathway were identified as positive regulators of the profibrotic response and genes in the ERK5 pathway as negative regulators. In contrast to these results, previous studies have reported that both JNK1/2 and ERK5 contribute to fibrosis^15–18^. Although our unbiased screen revealed the opposite regulation of ERK5 for fibrotic response, it might depend on the context of the screening assay. Protein phosphatase 2A (PP2A) is also a non-canonical pathway and inactivates TGF-β receptors through dephosphorylation^6,7^. Consistent with this molecular mechanism, two regulatory subunits of PP2A, Ppp2r3c and Ppp2r5b, suppressed the profibrotic response (Figure 2A). Tumor necrosis factor α (TNF-α) is a proinflammatory cytokine, but its role in fibrosis is controversial^19–21^. TGF-β and TNF-α have bilateral crosstalk, where TNF-α activates the non-canonical JNK pathway via TAK1 but inhibits the canonical pathway via Smad7^22,23^, resulting in synergistic and antagonistic effects, particularly in cancer pathogenesis^24^. In our screen, several factors in the TNF-α pathway promoted the profibrotic response to TGF-β treatment. Notch signaling is a cell-to-cell communication pathway that regulates cell fate determination during development and maintains tissue homeostasis in adult tissues. When Notch receptors on one cell bind to ligands on an adjacent cell, the Notch intracellular domain (NICD) is cleaved, translocates to the nucleus, and interacts with transcription factors to regulate gene expression^25^. Notch signaling is also involved in fibroblast proliferation, myofibroblast differentiation, and induction of contractile phenotype through synergistic interaction with TGF-β^26,27^. Consistently, our screen revealed the substantial impact of Notch signaling components on the profibrotic response.

**Figure 2.**
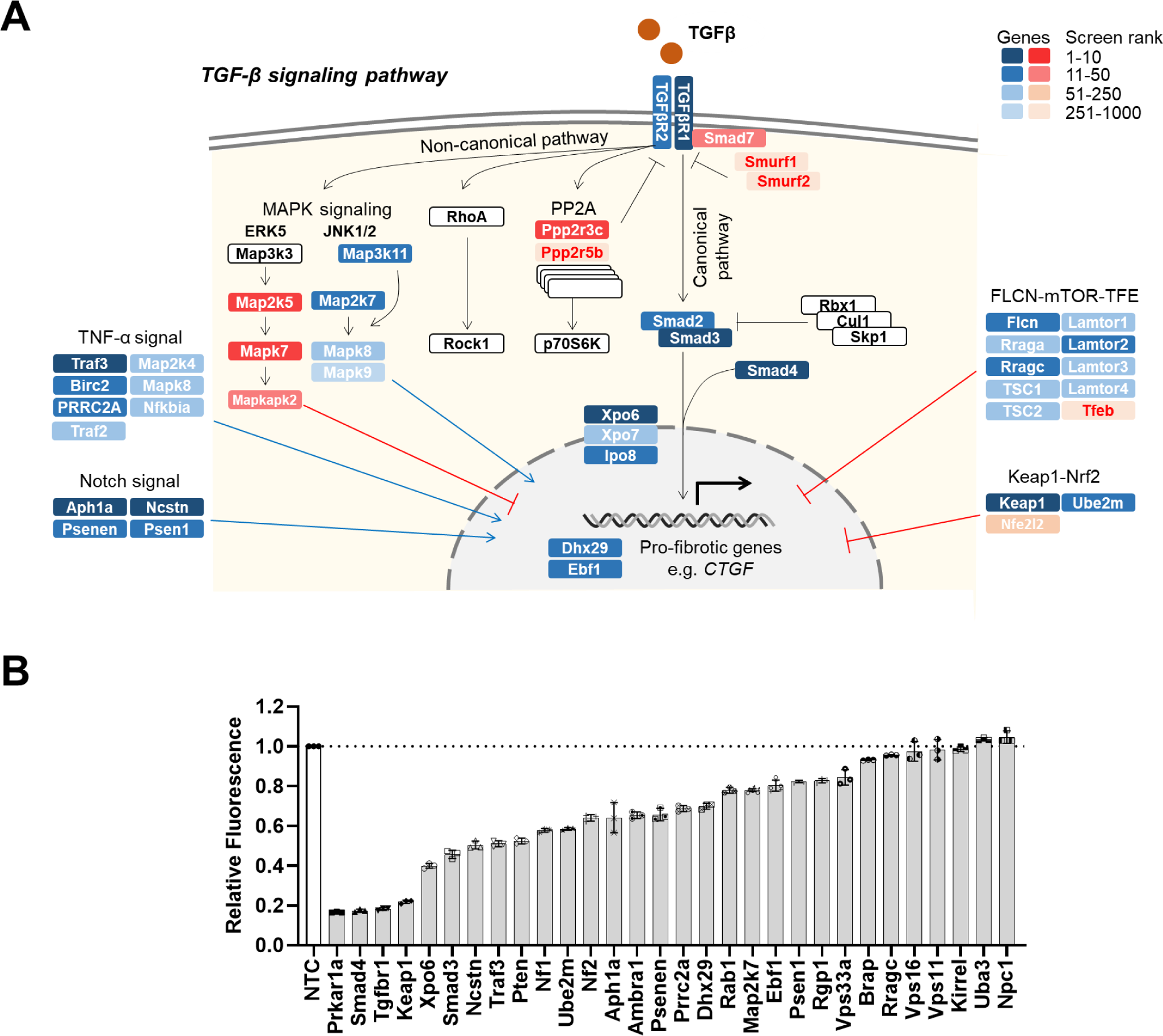
Individual gRNAs verify top hits from CRISPR screening. **A**, High-ranking genes were mapped to the TGF-β signaling pathway in the KEGG pathway database. **B**, Individual validation in the screening assay for the top 30 genes negatively regulating CTGF expression under TGF-β stimulation. Data are presented as mean±SD (n=3).

As a novel insight into pathways regulating TGF-β-mediated profibrotic response, clusters of Keap1-Nrf2 and FLCN-mTOR-TFE pathways were identified. Notably, Keap1 was the second top gene and individual gRNA experiments in the screening reporter assay confirmed the top impact on CTGF expression, similar to Smad4 and Tgfbr1 (Figure 2B). Thus, we focused on the Keap1-Nrf2 pathway and further investigated its impact on fibrosis and the molecular mechanism.

### Nrf2 suppresses the expression of fibrosis-related genes independently of antioxidant action

We generated the *Keap1* and *Nrf2* knockout C2C12 cells using CRISPR-Cas9 to verify the impact of the Keap1-Nrf2 pathway on fibrosis in response to TGF-β (Figure 3A). Endogenous expression of CTGF was reduced by *Keap1* knockout, which was reversed by additional *Nrf2* knockout, confirming that Nrf2 suppresses CTGF downstream of *Keap1* deletion. In addition to *Ctgf,* other fibrosis-associated genes such as *Postn*, *Acta2*, and *Fn1* were similarly regulated by the Keap1-Nrf2 pathway (Figure 3B). Next, we examined the anti-fibrotic effect of *Keap1* deletion in mouse embryonic fibroblasts (MEFs) using siRNA (Figure 3C). TGF-β treatment significantly increased the expression of CTGF and periostin, and *Keap1* knockdown abolished this upregulation. In addition, periostin expression was reduced below baseline, presumably due to spontaneous activation of fibroblasts in conventional cell culture (Figure 3D)^28^.

**Figure 3.**
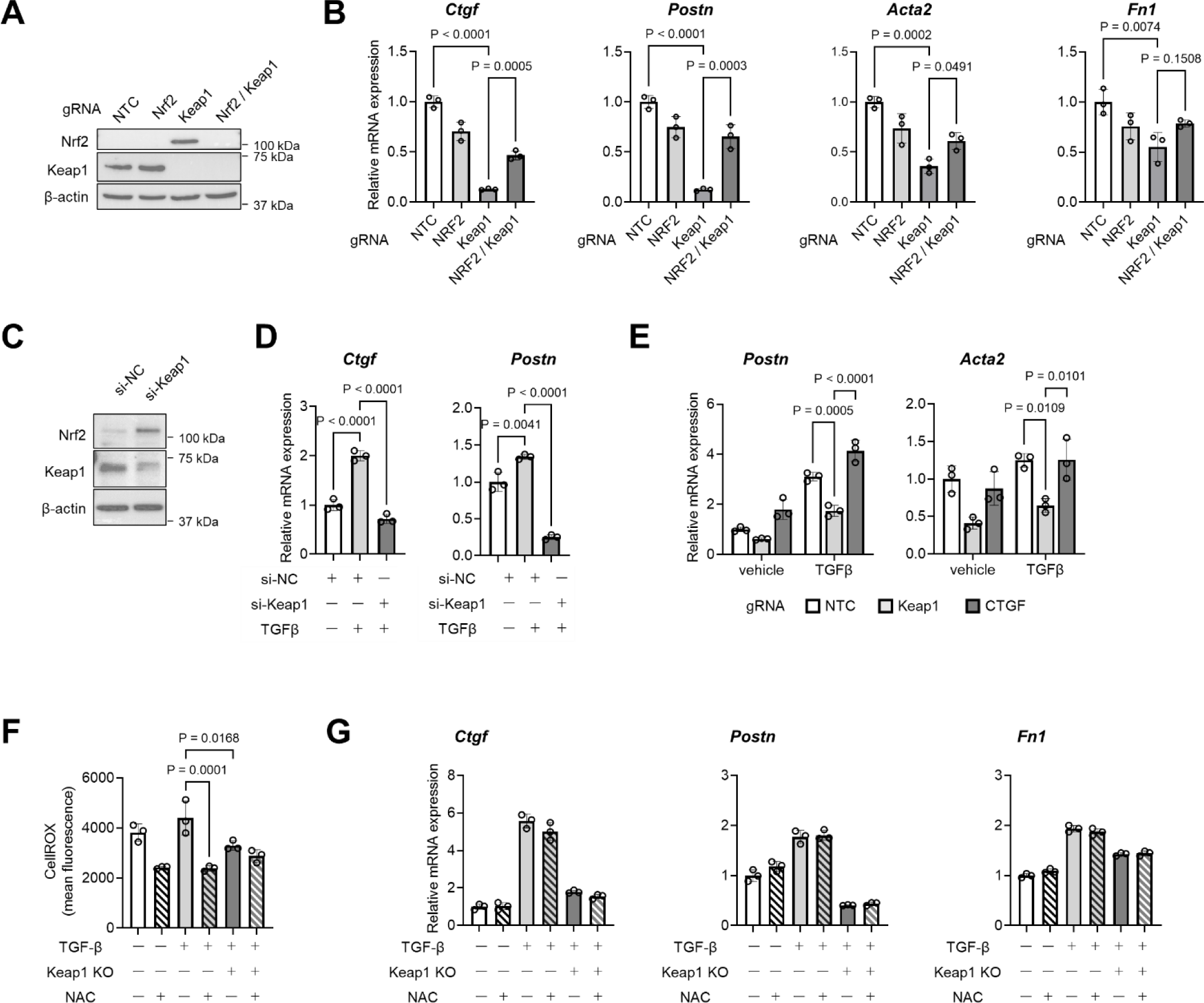
Nrf2 represses transcription of profibrotic genes independently of ROS regulation. **A**, Western blot to confirm knockout of *Keap1* and/or *Nrf2* in C2C12 cells. **B**, Quantitative PCR analysis of representative fibrosis-related genes in *Keap1* and/or *Nrf2* knockout C2C12 cells 24 hours after TGF-β treatment. **C**, Western blot to confirm the knockdown of *Keap1* and/or *Nrf2* in mouse embryonic fibroblasts (MEFs) using siRNA. **D**, Quantitative PCR analysis of *Ctgf* and *Postn* mRNA levels in *Keap1* and/or *Nrf2* knockdown MEFs 24 hours after TGF-β treatment. **E**, Quantitative PCR analysis of *Postn* and *Acta2* mRNA levels in *Keap1* or *Ctgf* knockout C2C12 cells at 24 hours after TGF-β treatment. **F**, Flow cytometry for CellRox to evaluate ROS production in Keap1 knockout C2C12 cells under 24-hour stimulation with TGF-β. N-acetylcysteine (NAC) treatment for 4 hours was employed as an antioxidant control. **G**, Quantitative PCR analysis of *Ctgf, Postn*, and *Fn1* mRNA levels in *Keap1* knockout C2C12 cells stimulated with TGF-β for 24 hours or NAC for 4 hours. Data are presented as mean±SD. *P* values were calculated by one-way ANOVA with Tukey’s multiple comparison test (n=3).

CTGF is a secreted glycoprotein that plays a central role in the fibrosis process. Genomic epidemiology studies have also shown that polymorphisms in the CTGF promoter region, which promotes CTGF transcription, are common in patients with systemic sclerosis^29^. CTGF is expected to be a therapeutic target for fibrotic diseases, and recently a phase II clinical trial demonstrated the therapeutic efficacy of the anti-CTGF monoclonal antibody, pamlevlumab, in preventing the progression of idiopathic pulmonary fibrosis^30^. When comparing the antifibrotic effect between CTGF and Keap1 inhibition, Keap1 knockout reduced profibrotic gene expression, but CTGF knockout showed no change, indicating that the Keap1-Nrf2 pathway has a broad fibrotic effect and may be a better therapeutic target than CTGF (Figure 3E).

Since the Keap1-Nrf2 pathway induces transcription of antioxidant genes, we investigated whether modulation of oxidative stress contributes to the antifibrotic response. TGF-β modestly increased reactive oxygen species (ROS) production, and *Keap1* knockout significantly reduced oxidative stress, as did N-acetylcysteine (NAC) treatment (Figure 3F). However, NAC treatment didn’t change the expression of profibrotic genes in baseline and TGF-β treatment with or without Keap1 knockout (Figure 3G). Taken together, these results indicated that when Keap1 was deleted, Nrf2 attenuated the profibrotic response to TGF-β treatment independently of the antioxidant effect.

### Nrf2 directly regulates the transcription of profibrotic genes

To investigate whether Nrf2 binds directly to profibrotic genes, we used Cleavage Under Targets and Tagmentation (CUT&Tag), which utilize a protein A-Tn5 transposase fusion protein to insert adapter sequences nearby the target protein. Amplification localized to the tagged regions provided efficient high-resolution sequencing libraries and higher sensitivity than conventional chromatin immunoprecipitation (ChIP) assays^31^. CUT&Tag was conducted with anti-Nrf2 antibody in Keap1 knockout cells. At profibrotic gene loci, Nrf2 binding was observed in the upstream regions of *Ctgf* and *Fn1* in C2C12 mesenchymal progenitor cells, whereas these Nrf2 binding peaks were not observed in mouse macrophage-like RAW264.7 macrophages. In contrast, Nrf2 binding near a representative antioxidant Nrf2 target gene locus, *Hmox1*, was observed in both C2C12 and RAW264.7 macrophages (Figure 4A). GO annotation analysis for genes enriched in C2C12 cells also showed that Nrf2 bound to gene loci associated with fibrosis-related gene sets, such as extracellular matrix (Figure 4B). Nrf2 negatively regulated the expression of profibrotic genes activated by the TGF-β-Smad pathway. Next, we examined the CUT&Tag with anti-Smad3 antibody in TGF-β treated wild-type and Keap1 knockout C2C12 cells. Smad3 binding was observed at similar loci upstream of *Ctgf* and *Fn1*, and these binding peaks were attenuated when Keap1 was deleted (Figure 4C). Smad3 binding was further confirmed by ChIP-qPCR analysis. In the peak regions proximal to *Ctgf* and *Fn1*, the Smad3 binding signal was increased upon TGF-β treatment, but was attenuated upon *Keap1* deletion (Figure 4D). Transcriptional inhibition of Nrf2 has been previously reported in the proinflammatory cytokine *IL6* and *IL1b* genes, where Nrf2 inhibits the recruitment of RNA polymerase II (Pol II) to the *IL6* and *IL1b* transcription start sites (TSSs)^9^. We also performed ChIP-qPCR analysis using anti-Pol II antibody to dissect the transcription initiation step. The peaks of Nrf2 and Smad3 binding were located at the proximity of TSSs of *Ctgf* and *Fn1* and Pol II binding to these sites was significantly reduced by Keap1 deletion (Figure 4E). Collectively, Nrf2 bound to loci upstream of *Ctgf* and *Fn1* and inhibited Smad3 binding and Pol II recruitment there, resulting in transcriptional inhibition of these genes.

**Figure 4.**
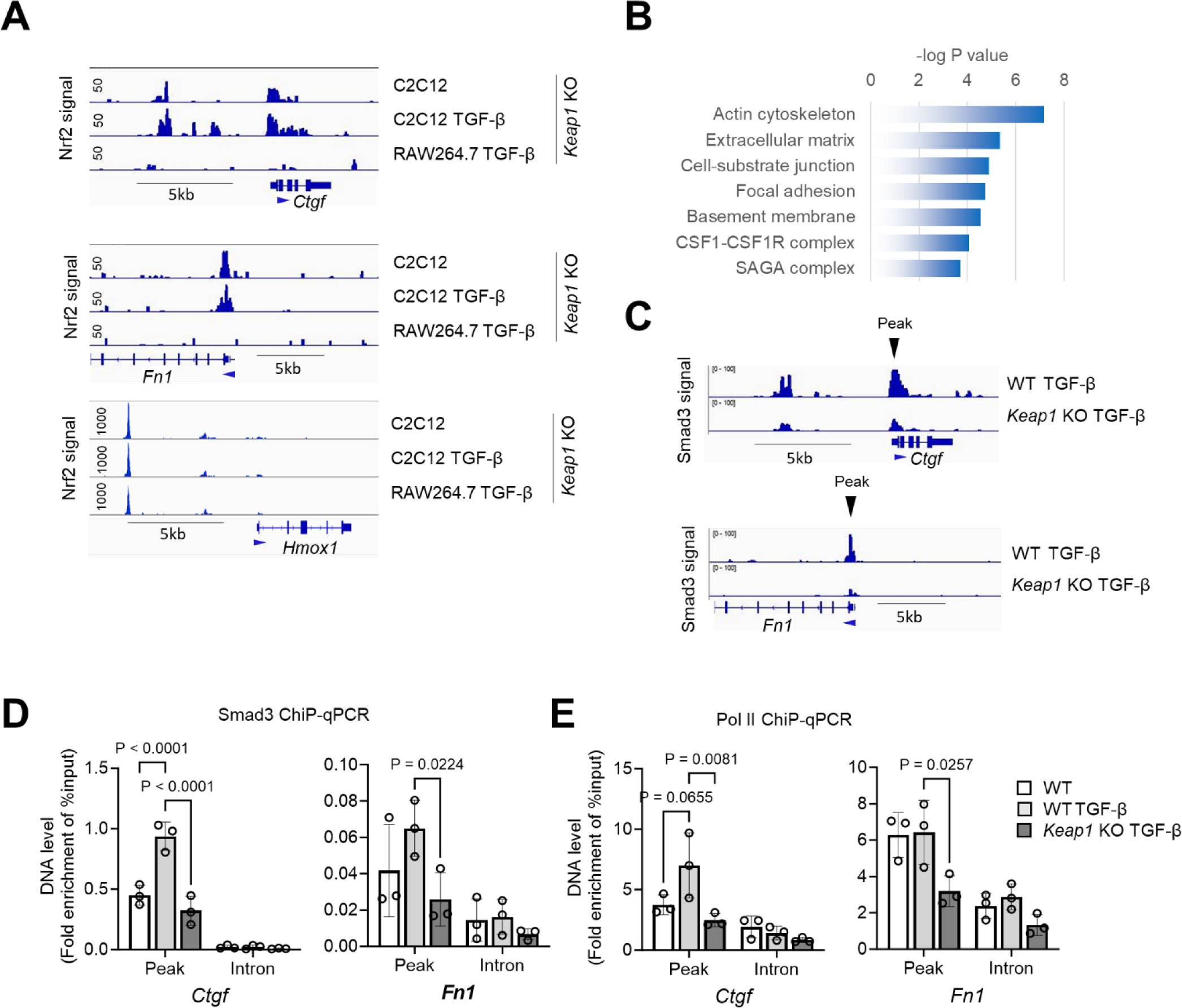
Nrf2 binds to the proximity of profibrotic genes and inhibits Smad3 binding and Pol II recruitment. **A**, Representative chromatin landscapes around *Ctgf*, *Fn1* and *Hmox1* regions of the mouse genome generated by CUT&Tag using anti-Nrf2 antibody in *Keap1* knockout C2C12 cells and *Keap1* knockout RAW264.7 macrophages with or without TGBF-β treatment. **B**, The result of GO annotation analysis. Nrf2-bound gene loci associated with fibrosis-related gene sets, such as extracellular matrix in C2C12 cells. **C**, Representative chromatin landscapes around *Ctgf* and *Fn1* regions of the mouse genome generated by CUT&Tag using anti-Smad3 antibody in C2C12 cells and *Keap1* knockout C2C12 cells with TBF-β treatment. **D**, *Smad3* ChIP-qPCR analysis of C2C12 cells or *Keap1* knockout C2C12 cells with or without TGB-β treatment. Primers were designed from the peak and intron regions of Smad3 binding. **E**, RNA polymerase II (Pol II) ChIP-qPCR analyses of C2C12 cells or Keap1 knockout C2C12 cells with or without TGB-β treatment. Primers were created from the peak and intron regions of Smad3 binding. Data are presented as mean±SD. *P* values were calculated by two-way ANOVA with Tukey’s multiple comparison test (n=3).

When we examined AREs proximal to profibrotic genes, we found no conserved TGAG/CnnnGC motif at the peak summits. However, in addition to known motifs, DNA-binding motif enrichment with HOMER analysis revealed several *de novo* motifs (Figure S2A). The second enrichment was the KLF5 motif and was observed in the peaks proximal to *Ctgf* and *Fn1* (Figure S2B). Typically, Nrf2 binds to AREs to activate the transcription of cytoprotective genes in a heterodimeric form with a group of small Maf proteins, such as MafF, MafG and MafK^32,33^. However, Nrf2 can also bind to non-AREs without small Maf^34^. Our motif analysis identified the candidate motifs for these atypical bindings. Similarly, no conserved TGAG/CnnnGC motif was observed in the transcriptional inhibition of proinflammatory cytokines, except IL6^9^. Altogether, it is suggested that Nrf2 binds to non-AREs without small Maf to repress the expression of profibrotic genes.

### Ddx54 contributes to Nrf2-mediated transcriptional inhibition of profibrotic genes

A corepressor is a protein that binds to transcription factors and represses gene expression. Identification of Nrf2-interacting partner proteins advances our understanding of the molecular mechanism underlying Nrf2-mediated transcriptional inhibition of profibrotic genes. We thus conducted a proximity-dependent biotin labeling technique to investigate the interactome of Nrf2. The biotin ligase fused to a target protein biotinylates proteins in close proximity to the target protein^35^. We employed TurboID, a highly efficient mutant of biotin ligase^36^, and generated the knock-in Nrf2-TurboID C2C12 cells to examine the physiological protein interaction. Similar to the *Ctgf-P2A-GFP* reporter cells, we obtained more than 10 clones of *Nrf2-HA-TurboID* cells (Figure S3A). Of these, clone 43, which yielded the same level of protein expression as wild-type Nrf2, was used in subsequent experiments (Figure S3B). Optimization of enzyme efficiency with labeling time and biotin concentration is important for low-abundance nuclear proteins. Therefore, we tested the effect of these parameters on biotin labeling. Weak background labeling was observed prior to incubation with biotin, which increased through 50 µM for 6 h and 500 µM for 10 min and saturated at 500 µM for 1 h and more (Figure S3C). As a control, we prepared tetracycline-inducible (Tet-On) *HA-TurboID-LNS* cells and titrated the concentration of doxycycline to balance the expression of TurboID. After the cells were treated with biotin at 500 µM for 10 min, biotinylated proteins were enriched with tamavidin 2-REV followed by LC-MS/MS analysis (Figure 5A). A total of 91 high confidence Nrf2 interactors were identified in this analysis (Table S1).

**Figure 5.**
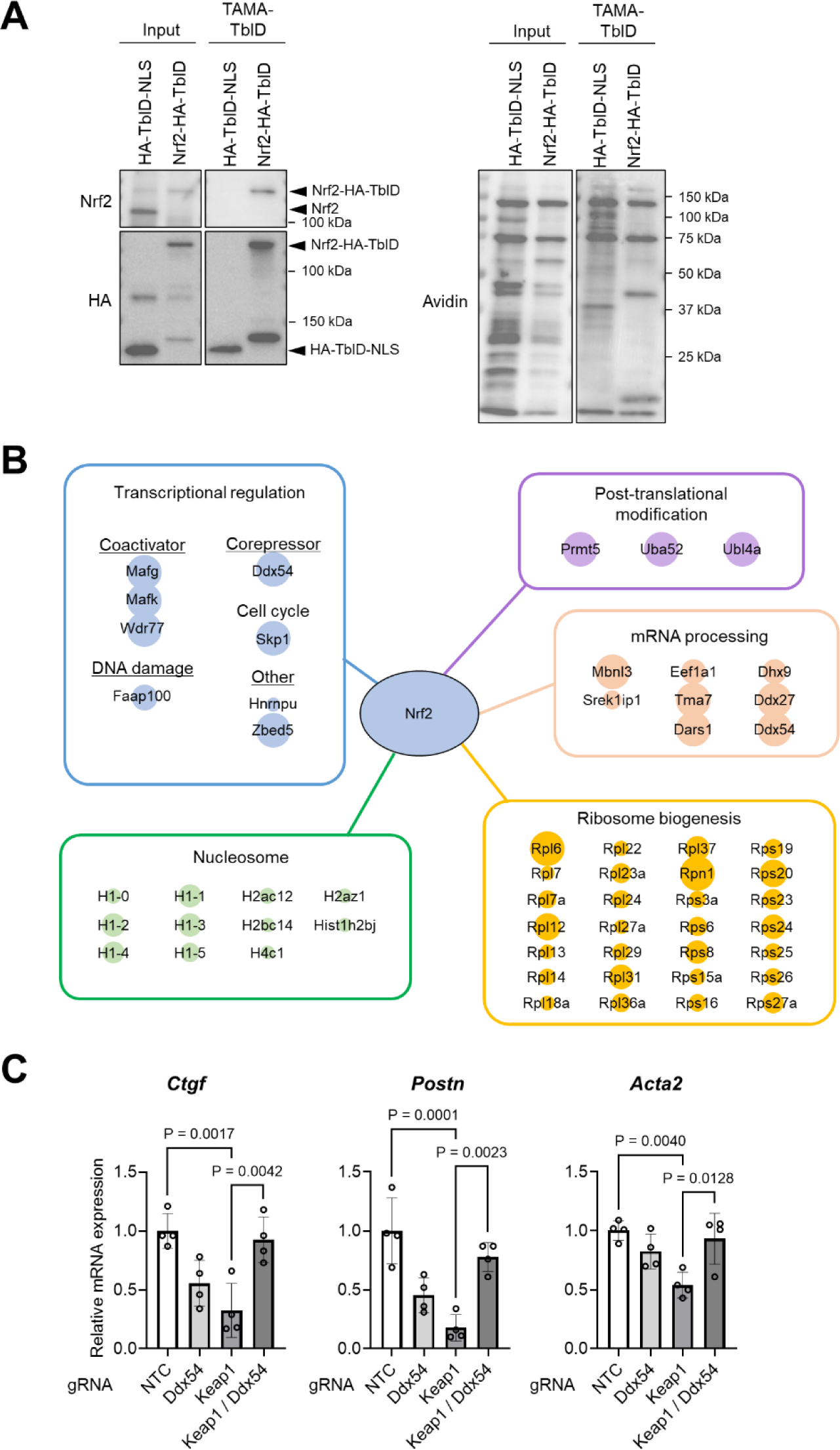
Ddx54 contributes to Nrf2-mediated transcriptional inhibition of profibrotic genes as a Nrf2-interacting corepressor. **A**, Western blot to confirm input and immunoprecipitated samples from *Nrf2-HA-TurboID* cells and (Tet-On) *HA-TurboID-NLS* cells used for BioID analysis. **B**, The result of BioID. 91 high-confidence Nrf2 interactors were identified. **C**, Quantitative PCR analysis of *Ctgf*, *Postn*, and *Acta2* mRNA levels in Keap1 and/or *Ddx54* knockout C2C12 cells 24 hours after TGF-β treatment. Data are presented as mean±SD. *P* values were calculated by one-way ANOVA with Tukey’s multiple comparison test (n=4).

The interactome was largely composed of nucleosome and ribosome component proteins, mRNA processing proteins, transcriptional regulators and their modifiers (Figure 5B). Canonical interacting partners, MafG and MafK, were identified, supporting the quality of this TurboID screen. Ddx54 was identified as a candidate repressor that interacts with Nrf2. Ddx54 is a member of the DEAD box proteins and has diverse cellular functions ranging from DNA transcription to RNA processing and translation^37^. Since Ddx54 has been reported to interact with estrogen receptors or splicing factor 1 and repress their transcriptional activity^38,39^, we investigated the contribution of Ddx54 to transcriptional inhibition by Nrf2. The transcription of the profibrotic genes *Ctgf*, *Postn,* and *Acta2* was repressed in *Keap1* knockout, but the additional knockout of *Ddx54* rescued these repressions, collectively indicating that Ddx54 interacts with Nrf2 to inhibit the transcription of profibrotic genes (Figure 5C).

### Loss of Keap1 in cardiac fibroblasts attenuates the hypertrophy and fibrosis

In order to examine the anti-fibrotic effect of Keap1 deletion in vivo, we used CRISPR/Cas9 to generate new alleles of Keap1 in which exon 3 is flanked by loxP sites (Figure S4A-B). Exons 3-6 encode the Kelch domain that binds to Nrf2, thus flanking exon 3 ensured loss of function through Cre-mediated recombination of the loxP sites. *Keap1^fl/fl^* mice were then crossed with *Postn^MCM^* mice^40^ to generate tamoxifen (TAM)-inducible fibroblast-specific *Keap1* knockout mice. Mice were subjected to transaortic constriction (TAC) surgery to induce pressure overload heart failure and compared to sham-operated controls (Figure S4C). Adult 9-week-old male mice were subjected to TAC or sham surgery and Cre activity was induced by intraperitoneal injection followed by feeding of TAM (Figure 6A). At 4 weeks post-TAC, control mice exhibited increased heart size and weight, indicative of cardiac hypertrophy, compared to sham-operated controls. However, these hypertrophic phenotypes were substantially reduced by *Keap1* deletion in the *Postn* lineage (Figure 6B-C). M-mode echocardiography was performed to assess cardiac function and dimensions. Contractile dysfunction and robust hypertrophy, as indicated by ejection fraction (EF), left ventricular (LV) mass and wall thickness, were observed in mice subjected to TAC-induced pressure overload for 4 weeks compared to sham-operated groups. However, *Postn^MCM^;Keap1^fl/fl^* mice showed significantly preserved LV size and function as indicated by improved EF and reduced left LV mass compared to *Postn^MCM^;Keap1^wt/wt^*controls (Figure 6D). Next, we evaluated the effects of *Keap1* deletion on cardiac fibrotic remodeling by visualizing collagen deposition in heart sections using Masson’s trichrome staining. TAC-operated hearts exhibited substantial fibrosis compared to sham-operated controls. However, deletion of *Keap1* in the *Postn* lineage significantly reduced the fibrotic area at 4 weeks after TAC (Figure 6E). Gene expression associated with fibrosis and heart failure was examined in the whole heart. TAC-induced pressure overload for 4 weeks robustly increased the mRNA expression of *Ctgf, Postn, Fn1, Cola1, Col3a1*, and *Col8a1*, but these upregulations were significantly attenuated by *Postn* lineage-specific Keap1 knockout. Similar transcriptional changes were observed in heart failure-related genes, including *Nppa, Nppb,* and *Myh7* (Figure S5).

**Figure 6.**
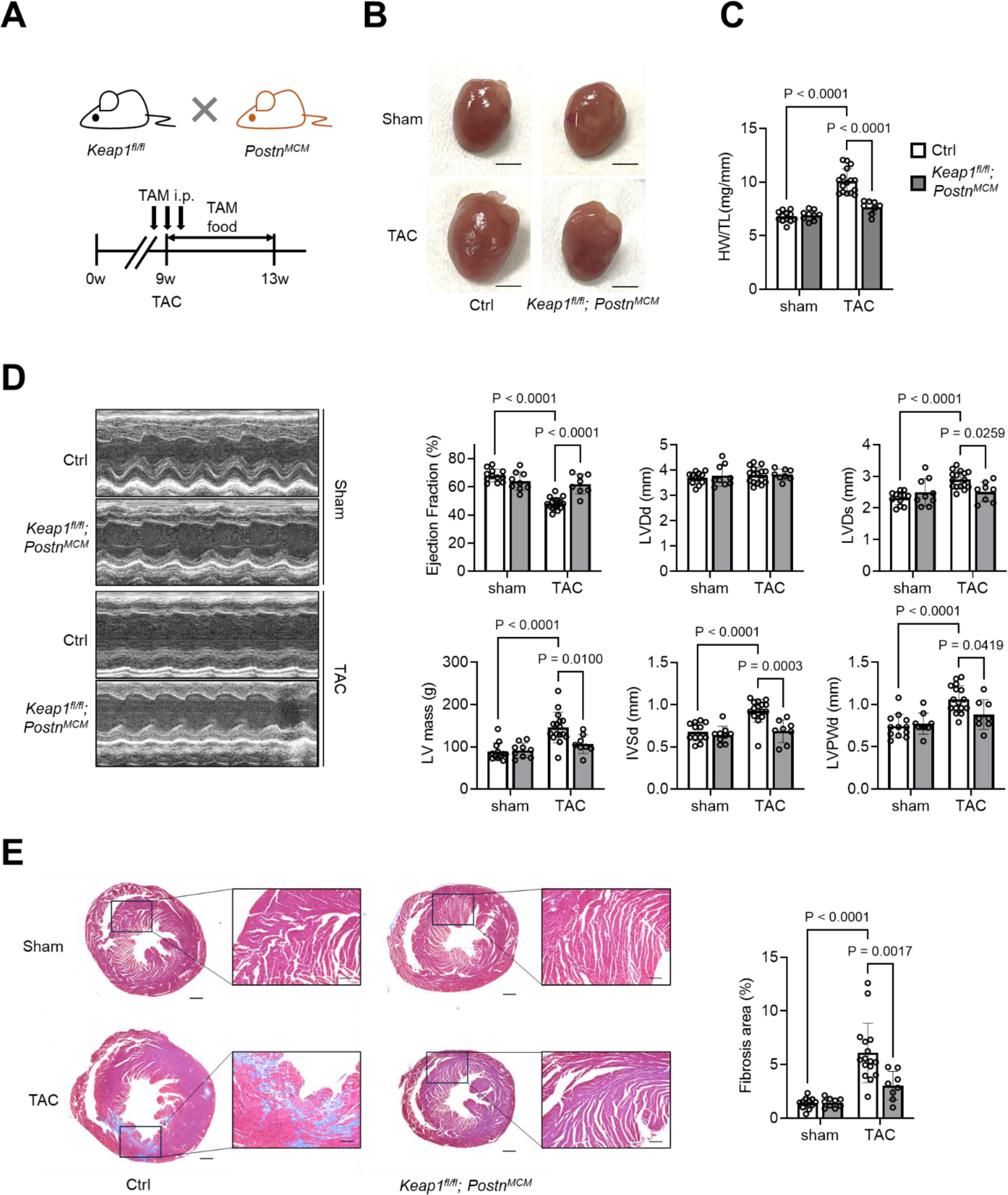
Loss of Keap1 in cardiac fibroblasts attenuates the heart hypertrophy and fibrosis. **A**, *Keap*1^fl/fl^ mice were crossed with *Postn-MerCreMer* (*Postn^MCM^*) mice to generate tamoxifen-inducible fibroblast-specific *Keap1* knockout mice. Adult 9-week-old male mice were subjected to trans-aortic constriction (TAC) surgery or sham surgery and Cre activity was induced by intraperitoneal injection followed by feeding with tamoxifen. Mice were sacrificed 4 weeks after TAC or sham surgery. **B**, Representative photographs of hearts from fibroblast-specific *Keap1*-deficient (*Keap1*^fl/fl^, *Postn^MCM^*) and control (*Postn^MCM^*) mice 4 weeks after TAC or sham surgery. Scale bars: 5 mm. **C**, Quantitative analysis of heart weight to tibial length ratio. **D**, Representative echocardiographic M-mode images of left ventricles and ejection fraction (EF), left ventricular (LV) mass and wall thickness. **E**, Representative photomicrographs of transverse sections stained with Masson’s trichrome and quantification of interstitial fibrosis area. Scale bars: 500 µm. Data are presented as mean±SD. *P* values were calculated by two-way ANOVA with Tukey’s multiple comparison test (n=8-16 each).

### Cardiomyocyte-specific deletion of Keap1 has no effect on the hypertrophy and fibrosis

Historically, Nrf2 has been considered a master regulator of antioxidant defense, and cardiomyocyte Nrf2 may also influence the progression of heart failure^41^. We thus investigated the effect of *Keap1* deletion in cardiomyocytes. *Keap1^fl/fl^* mice were crossed with *Myh6^MCM^* mice^42^ to generate TAM-inducible cardiomyocyte-specific Keap1 knockout mice. Cardiac phenotypes were then analyzed under pressure overload for 4 weeks (Figure S6A). TAC-operated mice exhibited cardiac hypertrophy, which was similar between control and *Myh6^MCM^;Keap1^fl/fl^* mice (Figure S6B-C). In M-mode echocardiography, EF, LV mass, and other dimensions were not different between the two groups (Figure S6D). Similarly, deletion of *Keap1* in the *Myh6* lineage did not affect collagen deposition in pressure overload heart failure (Figure S6E). In the context of TAC-induced pressure overload for 4 weeks, the Keap1-Nrf2 pathway in fibroblasts, but not in cardiomyocytes, modulated cardiac hypertrophy and fibrosis.

### Fibroblast-specific deletion of Keap1 results in attenuation of renal fibrosis

Finally, we employed the unilateral ureteral obstruction (UUO) model of progressive renal fibrosis^43^ to investigate the universal antifibrotic effect of *Keap1* deletion in the *Postn* lineage. Adult 7-week-old male mice were treated with TAM by intraperitoneal injection followed by chow, and then subjected to UUO or sham surgery at 9 weeks of age (Figure 7A). In response to UUO, sham-operated control mice exhibited atrophic morphology and decreased kidney weight. In contrast, *Keap1* deletion in the *Postn* lineage induced atrophy at baseline but no further weight loss in the UUO model (Figure 7B-C). On histological analysis with Masson’s trichrome staining, UUO-operated kidney showed robust interstitial collagen deposition and parenchymal thinning, whereas these changes were mild and fibrotic areas were reduced in UUO-operated *Postn^MCM^;Keap1^fl/fl^* mice (Figure 7D). Consistently, UUO surgery induced substantial upregulation of fibrosis-related gene expression, but *Keap1* deletion in the *Postn* lineage attenuated these profibrotic transcriptions (Figure 7E).

**Figure 7.**
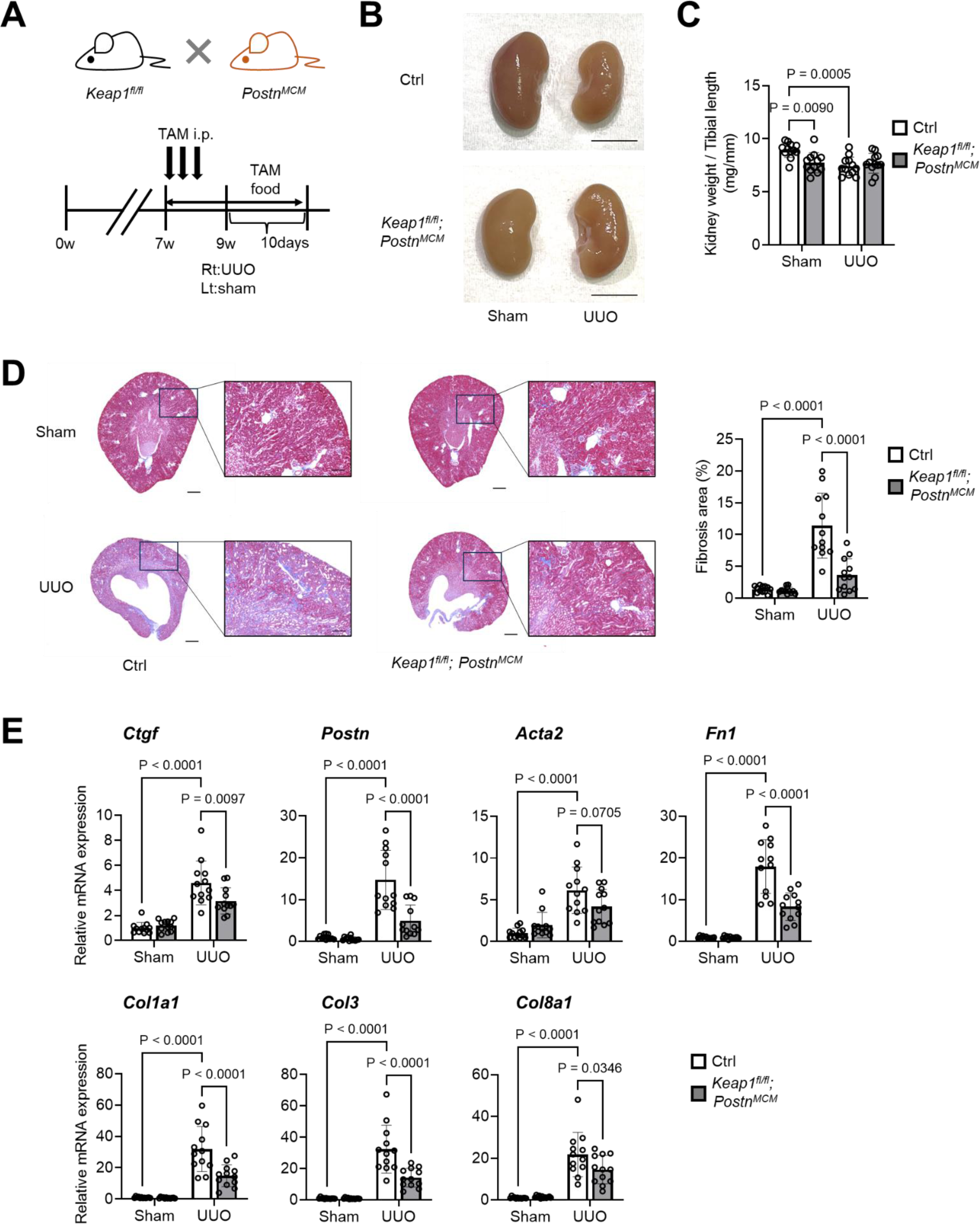
Loss of Keap1 in renal fibroblasts attenuates the renal atrophy and fibrosis. **A**, Adult 7-week-old male mice were treated with tamoxifen by intraperitoneal injection and subsequent feeding, and then subjected to UUO (right kidney) or sham surgery (left kidney) at 9 weeks of age. Mice were sacrificed 10 days after UUO surgery. **B**, Representative photographs of kidneys from fibroblast-specific *Keap1*-deficient (*Keap1^fl/fl^*, *Postn^MCM^*) and control (*Postn^MCM^*) mice 10 days after UUO or sham surgery. Scale bars: 5 mm. **C**, Quantitative analysis of the ratio of kidney weight to tibial length. **D**, Representative photomicrographs of Masson’s trichrome stain sagittal sections and quantification of interstitial fibrosis area of kidneys. Scale bars: 500µm. **E**, Quantitative PCR analysis of representative fibrosis-related genes in whole kidney tissue. Data are presented as mean±SD. *P* values were calculated by two-way ANOVA with Tukey’s multiple comparison test (n=12 each).

## Discussion

Activated fibroblasts are the central effector cells in organ fibrosis and serve as the primary ECM-producing cells. The transformation of fibroblasts into secretory, matrix-producing, and contractile myofibroblasts is an important cellular event in many fibrotic diseases. There is extensive crosstalk between myofibroblasts, innate and adaptive immune cells, vascular endothelial cells, and parenchymal cells via cytokines, growth factors, and matricellular proteins^44^. Of these, TGF-β is a prominent player in the initiation, progression, and persistence of fibrosis, primarily affecting fibrosis and epithelial cells^45^. Accumulating evidence supports the pathological role of TGF-β in various organ fibrosis diseases, including heart failure and chronic kidney disease. Despite this compelling evidence, the therapeutic translation of anti-TGF-β strategies in patients with fibrotic diseases has been challenging due to the broad biological effects of TGF-β and the complexity of fibrotic responses^46^. Based on this background, this study focused on fibroblast activation by TGF-β and identified the Keap1-Nrf2 pathway as a novel mechanism to inhibit fibrosis.

Although the antifibrotic effect of Nrf2 has been observed previously, it has been assumed that Nrf2 primarily suppresses oxidative stress via the expression of antioxidant genes such as GSTs, HO-1, and NQO1, and the consequent inhibition of ROS injury or signaling contributes to antifibrosis^47^. In contrast, we discovered that Nrf2 directly regulates the transcription of profibrotic genes independently of ROS scavenging, at least in the context of fibroblast activation. Classically, Nrf2 was thought to recognize AREs with small Maf and act as a transcriptional activator. In addition, recent paper indicated that Nrf2 could alternatively bind to sequences other than AREs and act as a transcriptional repressor in the regulation of proinflammatory cytokines^9^. It was a turning point in understanding the versatile gene regulation of Nrf2^41^, but the molecular basis of how Nrf2 elicits transcriptional inhibition remained to be elucidated. Similar to this anti-inflammatory regulation, the present study shows that NRF2 also negatively regulates the expression of profibrotic genes. As a mechanism, CUT&tag revealed that NRF2 binds to the promoter region and is associated with decreased binding of the profibrotic transcription factor Smad3 and Pol II. Additional DNA-binding motif enrichment analysis indicated that the KLF5 motif is the potential sequence for NRF2 interaction, but it is still unclear whether Nrf2 binds there directly or through association with other factors. As a further mechanism, proximity-dependent biotin labeling with TurboID revealed that Ddx54 interacts with Nrf2 and contributes to transcriptional repression in the context of the TGF-β-mediated profibrotic response. Ddx54 has diverse cellular functions ranging from DNA transcription to RNA processing and translation^37^. Since the *Keap1* deletion-mediated mRNA reduction in profibrotic genes was almost completely abrogated by *Ddx54* deletion (Figure 5C), it is likely that Ddx54 inhibits Pol II recruitment and acts as a transcriptional repressor in the proximity of the profibrotic genes.

CTGF is a matricellular protein and its expression is elevated in human fibrotic diseases in various organs^48^. CTGF is therefore considered a promising therapeutic target in fibrotic diseases, and indeed clinical trials of monoclonal antibodies targeting CTGF are underway in pulmonary fibrosis and other diseases^30,49^. In heart failure, lineage-specific knockout experiments have shown that CTGF secreted by fibroblasts but not myocardium promotes cardiac fibrosis^50,51^, and anti-CTGF antibodies effectively inhibit excessive fibrosis and promote tissue repair after myocardial infarction in animal models^48^. Consistently, Keap1 deletion in fibroblasts prevented cardiac fibrosis, but cardiomyocyte-specific knockout failed to show an antifibrotic effect. Furthermore, this study identified the Keap1-Nrf2 pathway as a major negative regulator of CTGF expression in the CRISPR screen, but this pathway affected a wide range of fibrosis-related genes, making it a potentially more impactful therapeutic target than CTGF.

In conclusion, we have identified the Keap1-Nrf2 pathway as a key regulator of fibrosis by unbiased screening of fibroblast activation. Although this pathway was previously thought to exert an antifibrotic effect through its antioxidant action, this study demonstrates for the first time that it directly represses the transcription of fibrotic-related genes in fibroblasts. Although the mechanism of transcriptional repression of Nrf2 has remained unclear, the present study suggests that Ddx54 is associated with Nrf2 as a transcriptional repressor. And the antifibrotic effects of Keap1 deletion were confirmed in mouse models of heart failure and chronic kidney disease, making it a promising target for future antifibrotic therapy.

## Author contributions

A.H., and T.N. designed the study; T.N. executed most experiments with help from A.H., N.I., M.C.A., Y.H., Y.U., and S.T.; Y.K., Y.O.,and D.M. analyzed CRISPR library screening and CUT & Tag; R.K., T.H., and I.H. generated *Keap1^fl/fl^*mouse. S.M. supervised the research; T.N., and A.H. wrote the manuscript; All authors discussed the results and commented on the manuscript.

## Methods

### Cell cultures

Mouse C2C12 mesenchymal progenitor cells (RIKEN BioResource Center, Tsukuba, Japan), mouse MEFs (RIKEN BioResource Center, Tsukuba, Japan), mouse RAW264.7 macrophages (RIKEN BioResource Center, Tsukuba, Japan) and Lenti-X 293T (Clontech) were cultured at 37°C in 5% CO2 in Dulbecco’s modified Eagle’s medium (DMEM, FUJIFILM) containing 10% FBS (fetal bovine serum, HyClone) and penicillin-streptomycin (100 U/ml, Thermo Fisher Scientific). To promote fibroblast activation, C2C12 cells and MEFs were cultured with 10 ng/mL TGF-β (PEPRO TECH, #100-21) for 24 hours. All cell lines were routinely tested negative for mycoplasma contamination.

### Plasmids

Individual gRNAs were cloned into lentiCRISPR v2 (Addgene #52961) or lentiGuide-Puro (Addgene #52963) and cDNAs were cloned into pLenti (Addgene #22255) or pMSCV (Clontech). Drug resistance genes in lentiGuide and pLenti were replaced by blasticidin S, neomycin or puromycin resistance genes. pMD2.G (Addgene #12259) and psPAX2 (Addgene #12260) were used for lentiviral packaging.

### Virus production

To produce lentiviruses, 6-well plates of 70% confluent Lenti-X 293T cells (Clontech) were transfected with 1.5 μg transfer vector, 0.5 μg pMD2.G and 1.0 μg psPAX2 using Fugene HD (Promega) according to the manufacturer’s instructions. The supernatant was collected after 48 hours and frozen at −80°C.

### Knock-in cell generation

Using CRISPR-Cas9 system, the inserted fragment (*P2A-GFP-loxP-pPGK-puro-loxP* or *HA-TurboID-loxP-pPGK-puro-loxP*) was inserted just before the stop codon of *Ctgf* or *Nrf2*, and the *pPGK-puro* cassette was removed by adenovirus-mediated Cre expression. The insertion was confirmed by PCR. The resulting knock-in cells were separated into single cells using a cell sorter (SH800S, Sony Imaging Products & Solutions Inc) and cultured. *Ctgf-P2A-GFP* knock-in cells were cultured with TGF-β and GFP enhancement was confirmed by flow cytometry. Western blot confirmed correct protein expression in *Nrf2-HA-TurboID* cells.

### Real-time quantitative PCR analysis

Total RNA was isolated from cells or heart and kidney tissues using TurboCapture mRNA Kits (QIAGEN). Please refer to the manufacturer’s instructions for details. Complementary DNA (cDNA) was synthesized from total RNA using PrimeScript RT Master Mix (TAKARA). Real-time quantitative PCR was performed with CFX384 Touch (Bio Rad) using KAPA SYBR FAST qPCR Master Mix (KAPABIOSYSTEMS).

### CRISPR Screening

Cas9-expressing *Ctgf-P2A-GFP* C2C12 cells were infected with validated lentiviral particles generated from a whole-genome CRISPR-Brie library (Addgene, #73632). After 24 hours, infected cells were treated with 5 μg/mL puromycin for 2 days. Cells were cultured for a further 10 days. The cells expressing the library of gene-specific gRNAs were then subjected to TGF-β treatment for 24 hours to activate the profibrotic response. The highest and lowest 20% of cells with GFP signal were collected by fluorescence-activated cell sorting (FACS) for genomic DNA isolation and deep sequencing.

### Deletion of target genes using CRISPR-Cas9 system

The expression vector for the expression of the guide RNA was generated using the LentiGuide-Puro plasmid (addgene, #52963). These plasmids were transfected into cells expressing Cas9. After 24 hours, we performed drug selection by puromycin stimulation and cultured for 14 days.

### Immunoblot

Lysates were prepared from subconfluent cells lysed in RIPA buffer (FUJIFILM) and centrifuged at 16,000g for 15 minutes. SDS sample buffer (BIO RAD) was added and samples were boiled for 5 minutes. Proteins were transferred to polyvinylidene difluoride (PVDF) membranes. Membranes were then probed with anti-Nrf2 (CST, 12721S, 1:1000 for WB), anti-Keap1 (MBL, M-224-3, 1:1000 for WB), anti-Smad3 (CST, 9523S, 1:1000 for WB), anti-Avidin HRP (Biolegend, 405103, 1:2500 for WB), anti-HA tag (CST, 2367, 1:1000 for WB) and anti-β-actin (Sigma, 2228, 1:2000 for WB) antibodies followed by incubation with HRP-conjugated secondary antibodies (CST, 7076S/7074S, 1:1000 for WB). Blots were visualized by ECL (Pierce ECL Western Blotting Substrate, Thermo Fisher Scientific,). Western blot images were acquired using the ChemiDoc Touch MP imaging system (Bio-Rad).

### Gene knockdown by siRNA

The role of Nrf2 signaling in MEF cells was investigated by inhibiting Keap1 using small interfering RNA (siRNA; siRNA mouse silencer select, Keap1 ID: s78526; Thermo Fisher Scientific). MEFs at 70% confluence were plated in 6-well plates 24 hours before transfection. The siRNA-liposomal complexes were prepared using Lipofectamine RNAiMAX Reagent (Thermo Fisher Scientific) according to the manufacturer’s instructions. The siRNA-liposomal complexes were added to MEFs and incubated for 24 hours. Medium was changed after 24 hours. Cells were harvested 48 hours after transfection.

### Analysis of Reactive Oxygen Species

Analysis of intracellular Reactive Oxygen Species (ROS) generation was performed by flow cytometry. Briefly, C2C12 or C2C12-Keap1 knockout cells were plated in 6-well plates and incubated for 24 hours. Cells were then treated with TGF-β (10 ng/mL, 24 hours) and NAC (1mM) or DMSO alone for 4 hours. Cells were harvested and stained with 500 nM CellROX Green Reagent (Thermo Fisher Scientific) for 60 minutes at 37°C and analyzed by flow cytometry. The fluorescence signal proportional to cellular ROS levels was excited by a laser at 488 nm and quantified on an SH800S (SH800S, Sony Imaging Products & Solutions Inc).

### CUT&Tag experiments

Cells treated with TGF-β (10 ng/mL, 24 hours) were harvested and centrifuged at 600g at RT for 3 minutes. Cells (500,000 cells/sample) were washed twice in 1.5 mL Wash Buffer (20 mM HEPES pH 7.5; 150 mM NaCl; 0.5 mM Spermidine; 1× Protease inhibitor cocktail). Concanavalin A coated magnetic beads (Bangs Laboratories) were prepared as described and 10 µL of activated beads were added per sample and incubated at RT for 15 min. 10 µL of Concanavalin A coated magnetic beads (Bangs Laboratories) were added per sample and incubated at RT for 10 min. After a quick spin to remove liquid, the bead-bound cells were resuspended in 50 µL Antibody Buffer (20 mM HEPES pH 7.5; 150 mM NaCl; 0.5 mM Spermidine; 1× Protease inhibitor cocktail; 0.05% Digitonin; 2 mM EDTA; 0.1% BSA) containing a 1:50 dilution of the appropriate primary antibody; mouse anti-Keap1 antibody (M-224-3, MBL) and rabbit anti-Smad3 antibody (9523S, Cell Signaling). Incubation was performed overnight at 4°C. After a quick spin to remove liquid, an appropriate secondary antibody was diluted 1:50 in 100 µL Dig-Wash buffer (20 mM HEPES pH 7.5; 150 mM NaCl; 0.5 mM Spermidine; 1× Protease inhibitor cocktail; 0.05% Digitonin) and cells were incubated at RT for 60 minutes. Samples were washed three times in 1 mL Dig-Wash buffer by gentle vortex to remove unbound antibodies. The 1:400 dilution of the pA-Tn5 adapter complex was prepared in Dig-300 Buffer (0.01% Digitonin, 20 mM HEPES, pH 7.5, 300 mM NaCl, 0.5 mM Spermidine, 1× Protease inhibitor cocktail). After removal of the liquid from the magnetic stand, 50-100 µL was added to the cells, and incubated with pA-Tn5 overnight at 4°C. The samples were washed three times in 1 mL Dig-300 buffer by gentle vortex to remove unbound pA-Tn5 protein. The cells were then resuspended in 300 µL tagmentation buffer (10 mM MgCl2 in Dig-300 buffer) and incubated at 37°C for 60 minutes in a water bath. To stop tagmentation, 10 µL of 0.5 M EDTA, 3 µL of 10% SDS and 2.5 µL of 20 mg/mL Proteinase K were added to each sample, and the samples were incubated at 55°C for 30 minutes. To extract the DNA, 300 µL Phenol/Chloroform/Isoamyl alcohol (NIPPON GENE, 311-90151) was added to each tube with vortex and the DNA layer was collected. DNA was precipitated with Ethachinmate (NIPPON GENE) and dissolved in 25 mL 1/10 TE buffer (1mM Tris-HCL pH 8.0; 0.1mM EDTA). To amplify the libraries, 21 µL of DNA was mixed with 2 µL of a universal i5 and a barcoded i7 primer, using a different i7 primer for each sample. 25 µL NEBNext HiFi 2× PCR Master Mix (New England BioLabs) was added to the DNA mixture. PCR was performed on this sample in a thermocycler using the following protocol: 72 °C for 5 min (gap filling); 98 °C for 30 s; 14 cycles of 98 °C for 10 s and 63 °C for 10 s; final extension at 72 °C for 1 min and hold at 8 °C.

### Proximity-dependent biotin labeling with TurboID

TurboID was performed according to the previously paper^52^. Briefly, we generated *Nrf2-HA-TurboID* knock-in cells using CRISPR-Cas9 system. As a negative control, we also generated doxycycline inducible *HA-TurboID-NLS* C2C12 cell using pCW57.1 plasmid. Five plates of 15 cm *Nrf2-HA-TurboID* cells and *HA-TurboID-NLS* cells were grown to 80% confluence. These cells were treated with CDDO-Im (10 nM, 48 hours; R&D SYSTEMS), TGF-β (10 ng/mL, 24 hours; PEPRO TECH), doxycycline (0.03 µg/mL, 24 hours; TAKARA) and Biotin (500 µM, 1 hour; Sigma-Aldrich). Cells were harvested by scraping and washed three times with 5 mL PBS. Cell pellets were lysed in 6 mL RIPA buffer (FUJIFILM) containing 1 mM EDTA and 1:100 protease inhibitor cocktail (Sigma-Aldrich). The lysate was rotated at 4°C for 30 min, sonicated 3 × 30 s (power: middle, Bioruptor Plus, Diagenode) and then centrifuged at 16000 g for 15 min at 4°C. 75 µL of lysate was frozen as input sample and the rest was used as sample for IP. Biotinylated proteins were isolated by affinity purification using 150 µL of washed MagCapture Tamavidin2-REV (FUJIFILM, #136-18341) with rotation overnight at 4°C. The beads were then washed with 7×1 mL of 50 mM ammonium bicarbonate (pH 8.0) containing 1:100 protease inhibitor cocktail. The 4/5 beads were sampled for MS/MS analysis and the remaining 1/5 beads were sampled for IP. SDS sample buffer (BIO RAD) and RIPA buffer (FUJIFILM) were added to the IP beads and boiled for 5 min. The supernatant was collected using a magnetic stand. MS/MS analysis was performed by CERI (Chemicals Evaluation and Research Institute, Japan). Proteins of IP and input samples were evaluated by Western blotting. The following antibodies; mouse anti-HA tag (Cell Signaling Technology, #2367, 1/1000), rabbit anti-Nrf2 (Cell Signaling, #12721, 1/1000), mouse anti-β-actin (Sigma Aldrich, #2228, 1/2000) and HRP Avidin (Biolegend, #405103, 1/2500) were used to confirm the corresponding bands.

### Chromatin immunoprecipitation

Cells treated with TGF-β were fixed with 1% formaldehyde and then quenched with 1.25 M glycine for 5 min at room temperature. After washing with cold PBS, the fixed samples were resuspended in RIPA buffer (FUJIFILM) containing 1 mM PMSF and protease inhibitor cocktail (Sigma-Aldrich). Samples were centrifuged at 1,000g for 10 minutes at 4°C. After a quick spin to remove liquid, nuclear pellets were resuspended in 250 µL NUC buffer (15 mM Hepes pH 7.5, 60 mM KCl, 15 mM NaCl, 0.32 mM sucrose, 3 μM CaCl2) and digested with 1/100 MNase (New England Biolabs). After addition of 250 µL sonication buffer (90 mM Hepes pH 7.9, 220 mM NaCl, 10 mM EDTA, 1% NP-40, 0.2% sodium deoxycholate, 0.2% SDS), nuclei were sonicated in ice water for 5×30 s (power: middle, Bioruptor Plus, Diagenode). Samples were centrifuged for 10 min, 13000 g at 4°C. Supernatants were collected and antibodies against Pol II (Santa, Cruz 1/24) and Smad3 (#9523S, Cell Signaling, 1/50) were added to the supernatants and the lysates were rotated at 4 °C overnight. 30 µL Dynabeads protein A (Thermo Fisher Scientific) were added to the lysates and rotated at 4°C for 2 hours. Using a magnetic stand, the beads were washed in 1 mL Low Salt buffer (20 mM Tris-HCL (pH 8.0), 2 mM EDTA, 0.1% SDS, 1% TritonX-100, 150 mM NaCl, 1 mM PMSF), 1 mL High Salt buffer (20 mM Tris-HCL (pH 8.0), 2 mM EDTA, 0.1% SDS, 1% TritonX-100, 500 mM NaCl, 1 mM PMSF), 1 mL LiCl buffer (10 mM Tris-HCL (pH 8.0), 1 mM EDTA, 0.25 M LiCl, 1% NP40, 1% sodium deoxycholate, 1 mM PMSF) and 1 mL TE buffer (10 mM Tris-HCL (pH 8.0), 1 mM EDTA). After removing the liquid with the magnetic stand, 400 µL Elution buffer (0.1 M NaHCO3, 1% SDS) and 16 µL 5 M NaCl were added to the beads and incubated overnight at 65°C. Then 8 µL RNase, 8 µL 0.5 M EDTA, 16 µL 1M Tris-HCL (pH 6.5) and 2 µL 20 mg/mL Proteinase K were added to the samples and rotated at 55°C for 2 hours. Lysates were retrieved using the magnetic stand. For DNA extraction, Phenol/Chloroform/Isoamyl alcohol (NIPPON GENE, 311-90151) was added to the lysates and the DNA layer was collected. DNA was precipitated with Ethachinmate (NIPPON GENE, 312-01791) and dissolved in TE buffer (10 mM Tris-HCL (pH 8.0), 1 mM EDTA). For ChIP-qPCR analyses, the % Input values of the immunoprecipitated samples were normalized to the input DNA. Primers were designed from the CUT&Tag results.

### Mice

*Keap1*-*flox* mice were generated at the Gunma University. To exclude exons 3, loxP was inserted before and behind exons 3 using the CRIPSR Cas9 system (gRNA: Lt: GTGCCTTTACTAGCTCAGAA, Rt: TGCCTTCATATGAGGTTGAG). Sequential electroporation was performed on 266 embryos as previously described^53^. 40 mice were born and genotyped by RFLP. As a result, we obtained four candidates. TA cloning of the flox region was performed and sequencing of the flox clone was confirmed. As a result, 3 mice with the desired flox allele were obtained. *Postn*-MerCreMer mice and *Myh6*-MerCreMer mice were purchased from THE JACKSON LABORATORY JAPAN, INC. C57BL/6J mice were purchased from SHIMIZU Laboratory Supplies Co., Ltd. All mouse experiments were approved by the Animal Care and Use Committee of Kyoto Prefectural University of Medicine. Mice were maintained in a specific pathogen-free animal facility under a 13:11 h light/dark cycle at an ambient temperature of 21°C. Age-and sex-matched mice were used for all animal experiments.

### Tamoxifen administration

For Cre expression, mice were treated with tamoxifen. Tamoxifen (Cayman Chemical) was dissolved in peanut oil at a concentration of 10 mg/ml by shaking for 2 hours at 37°C. After dissolution, the tamoxifen was stored at 4°C. The injection dose was determined on the basis of body weight (approximately 120 mg tamoxifen/kg body weight). The tamoxifen/peanut oil solution was administered intraperitoneally to the mice three times every other day. The mice were also given a tamoxifen-containing diet. The tamoxifen-containing diet was manufactured by ORIENTAL YEAST CO., LTD (Tokyo, Japan). Based on NIH-07PLD, 0.04% tamoxifen (Tokyo Chemical Industry, T2510) and 4.95% sucrose were added.

### TAC operation

TAC was performed to induce pressure overload in the mouse heart. 9-week-old male mice were anesthetized with 2% isoflurane inhalation. The anesthetized mice were then placed in the supine position on a heating pad (37°C) and intubated with a 20-gauge tube using a small animal respirator (SN-480-7, Shinano Seisakusho, Tokyo, Japan) at 120 breaths per minute. And 1.0-2.0% isoflurane inhalation was added to maintain anesthesia. Hair on the chest and abdomen was removed with a depilatory agent. The operation was performed under a microscope. The chest was opened at the second intercostal space and the aorta was exposed. After isolation of the aortic arch between the innominate and left common carotid arteries, the transverse aortic arch was ligated (7-0 silk string) with an overlying 28-gauge needle. After ligation, the needle was quickly removed and the skin closed. The sham operation was identical except that the string was not ligated.

### UUO surgery

Mice were anesthetized with isoflurane. The anesthetized mice were then placed in the prone position. The right ureter was exposed and ligated in two places with 4-0 silk. No ligature was performed in the sham-operated group. 10 days after surgery, the mice were sacrificed and the kidneys were harvested.

### Analysis of interstitial fibrosis

To detect collagen content, Masson’s trichrome staining was performed on heart and kidney sections by Applied Medical Research Laboratory (Osaka, JAPAN). Samples were photographed and analyzed using BZ-X810 (KEYENCE JAPAN). Interstitial fibrosis was determined as the percentage of collagen-stained area/total heart area.

### Statistical analyses

All data are presented as mean±SD. *P* values were calculated by two-sided paired *t*-tests, one-way or two-way ANOVA with Tukey’s multiple comparison test using GraphPad Prism software version 9. No statistical methods were used to pre-determine sample size. Sample size was based on experimental feasibility and sample availability. Samples were processed in random order.

---

**Figure S1.**
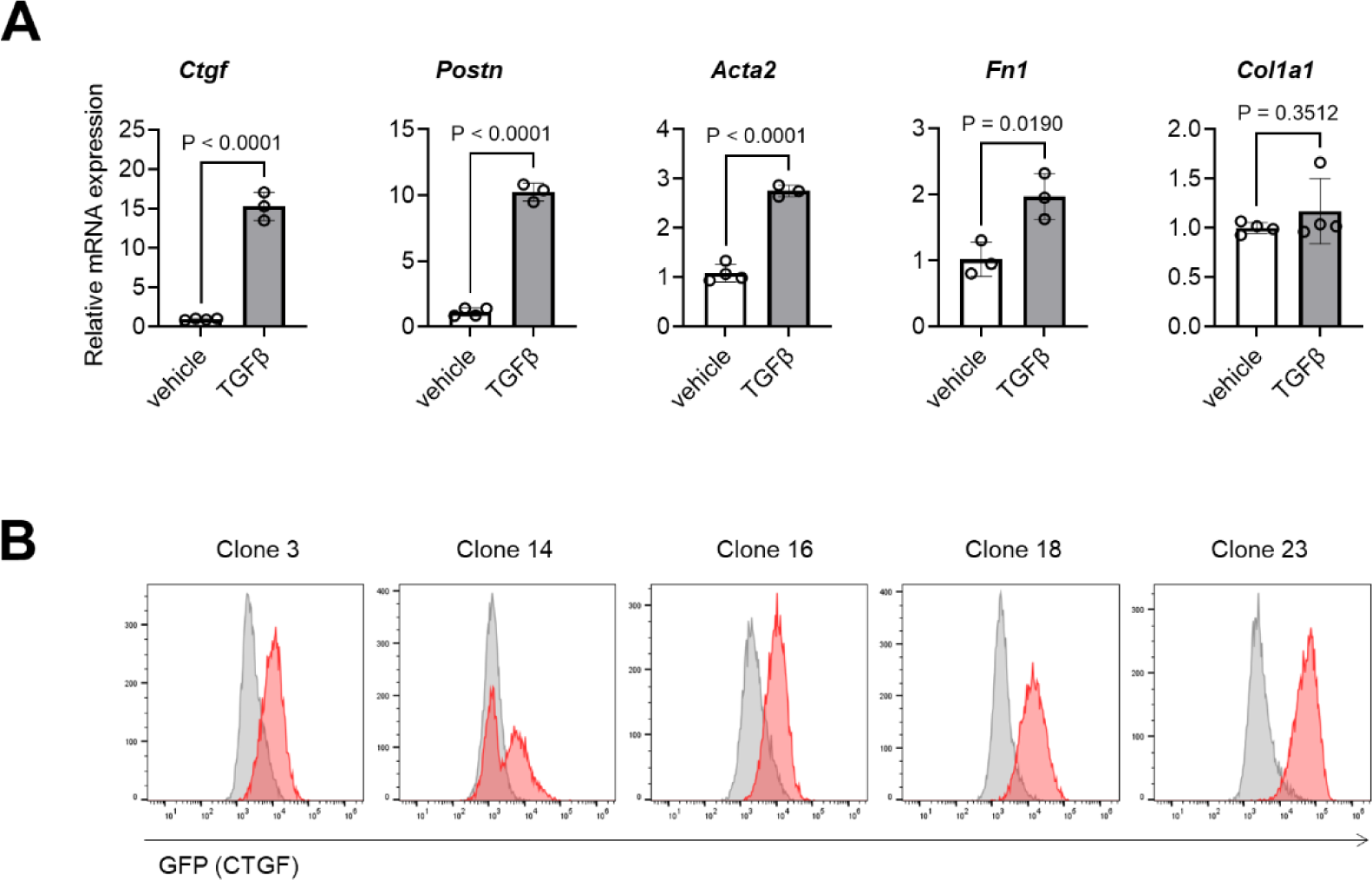
CTGF expression is robustly enhanced by TGF-β treatment and is reflected as a GFP signal in *CTGF-P2A-GFP* knock-in reporter cells. A, Quantitative PCR analysis of representative fibrosis-related genes in C2C12 mesenchymal progenitor cells 24 hours after TGF-β treatment. **B**, The response of the five clones of *CTGF-P2A-GFP* reporter cells to TGF-β treatment was evaluated by flow cytometry. Data are presented as mean±SD. *P*-values were determined by two-sided paired *t*-tests (n=3-4).

**Figure S2.**
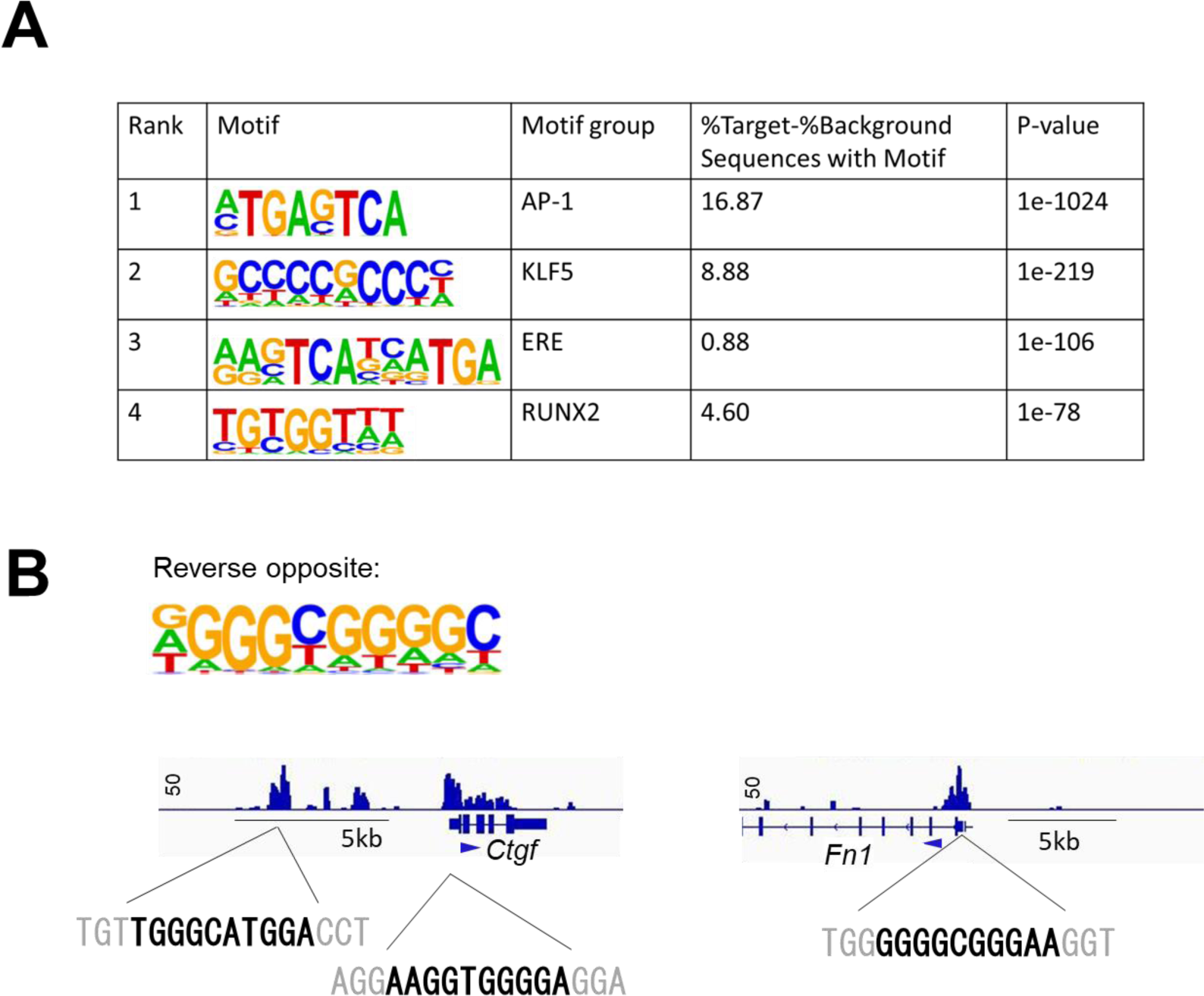
The analysis of the binding motifs of Nrf2. **A**, The *de novo Nrf2* binding motifs revealed by HOMER analysis. **B**, The KLF5 motif was observed in the peaks proximal to *Ctgf* and *Fn1*.

**Figure S3.**
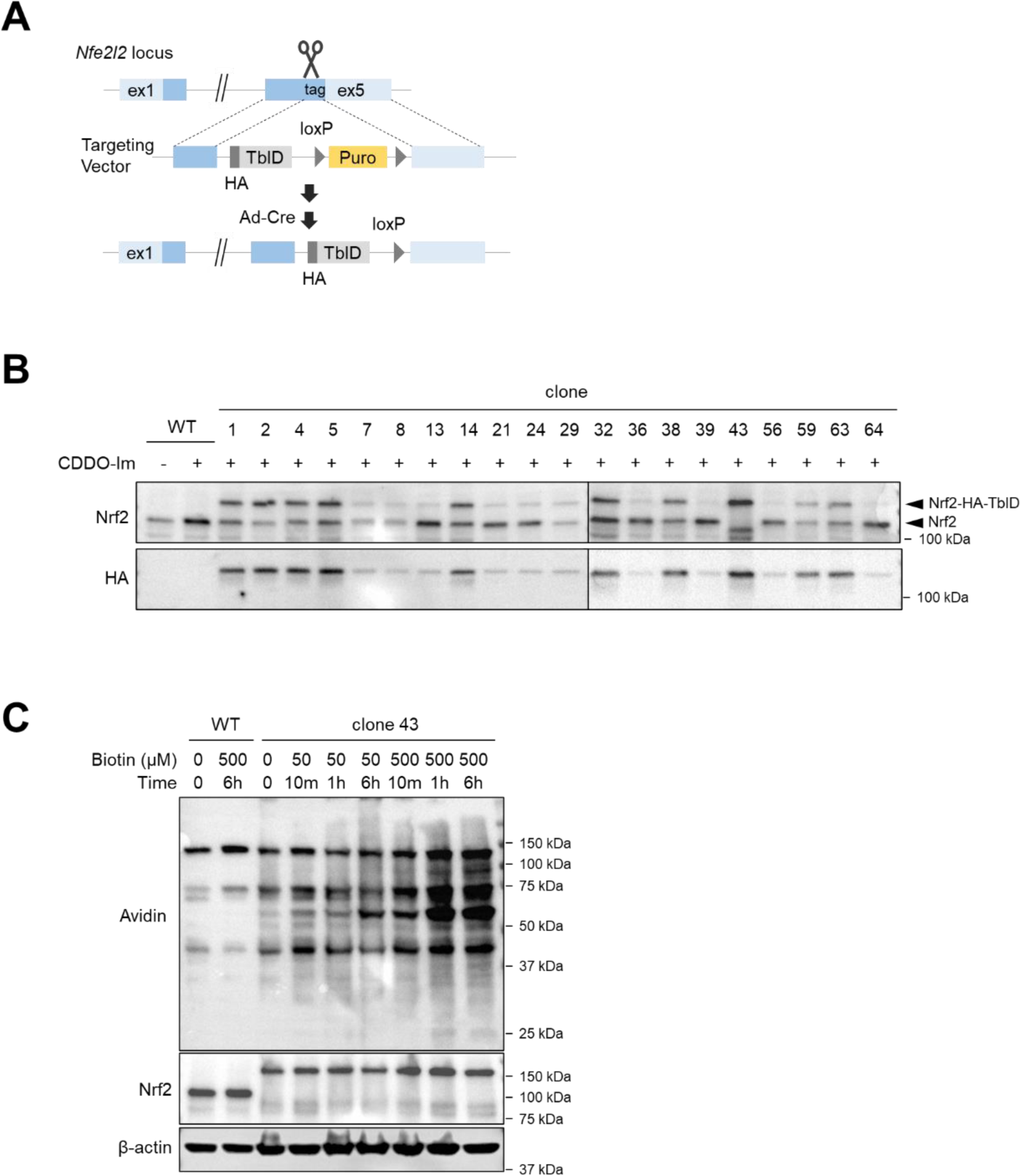
Generation of Nrf2-HA-TurboID knock-in cells and optimization of enzyme efficiency. **A**, The *Nrf2-HA-TurboID* knock-in cells were generated with mouse C2C12 mesenchymal progenitor cells using the CRISPR-Cas9 system. The *P2A-HA-TurboID-loxP-pPGK-puro-loxP* fragment was inserted just before the stop codon of *Nrf2* gene, and the *pPGK-puro* cassette was removed by adenovirus-mediated Cre expression. **B**, Western blot to confirm the knock-in clones after 24-hour treatment with Nrf2-activating CDDO-IM (10 nM). **C**, Western blot to confirm clone 43 of *Nrf2-HA-TurboID* cells for optimization of enzyme efficiency with labeling time and biotin concentration.

**Figure S4.**
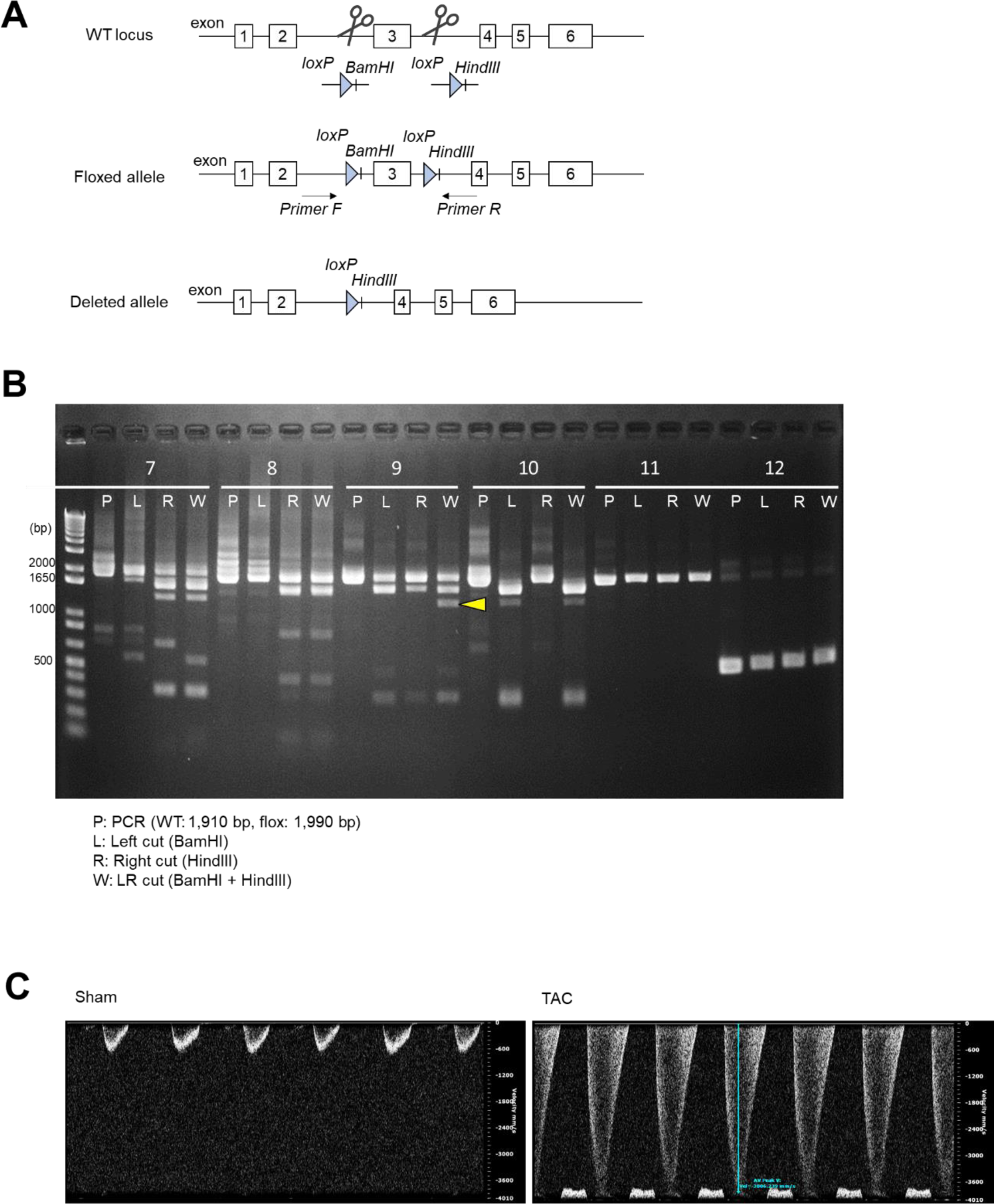
Generation of *Keap1* flox mice and TAC surgery. **A**, Generation of a new allele of *Keap1* in which exon 3 are flanked by loxP sites. Cre-mediated recombination of the loxP sites resulted in deletion of exon 3. **B**, PCR evaluation of genotyping for exon 3 deletion in candidate mice. **C**, Representative Pulsed Wave Doppler images from echocardiography 4 weeks after sham or TAC surgery.

**Figure S5.**
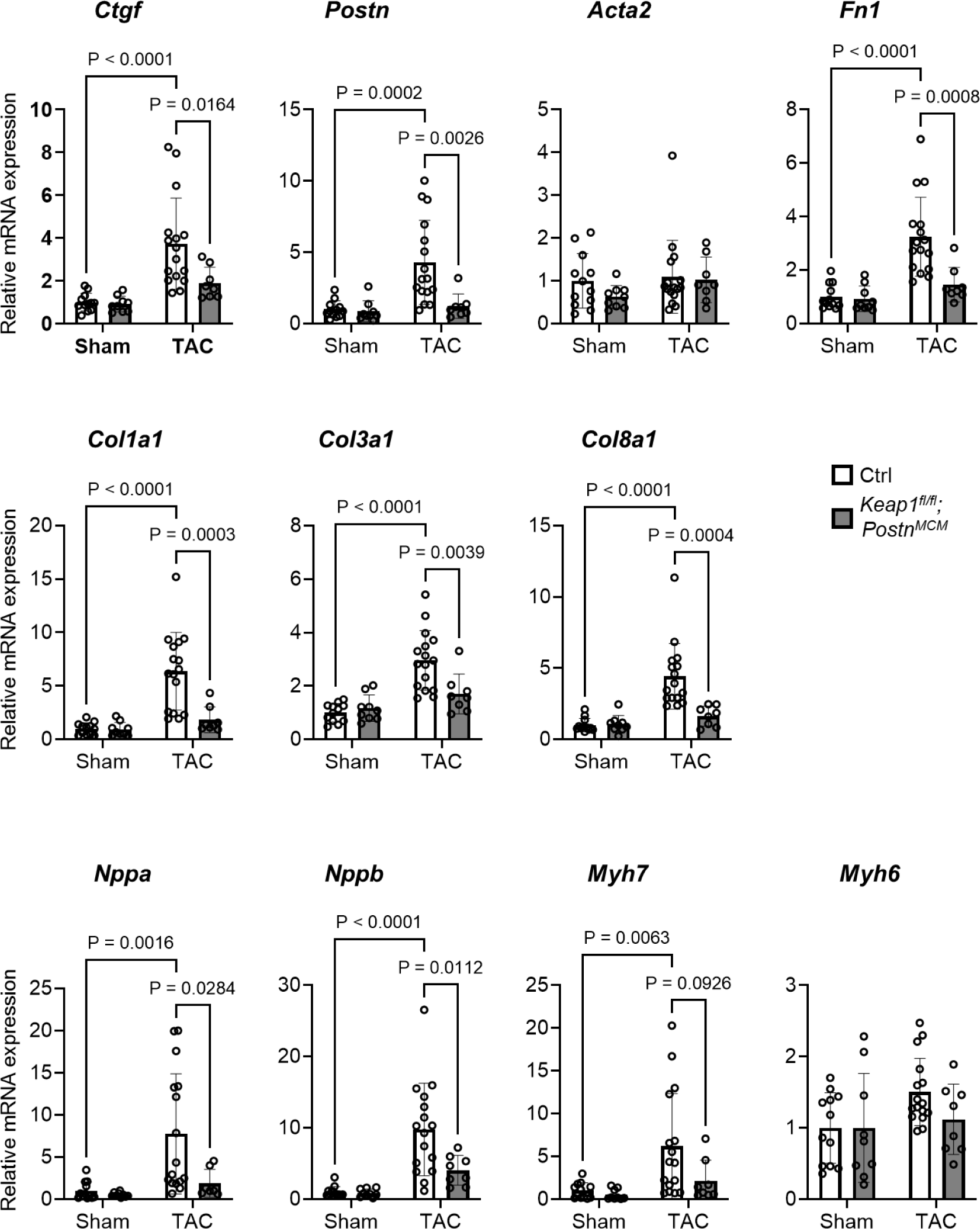
*Keap1* knockout in fibroblasts attenuates the gene expression associated with fibrosis and heart failure. Quantitative PCR analysis of genes associated with fibrosis and heart failure in the whole hearts of fibroblast-specific *Keap1*-deficient (*Keap1^fl/fl^*, *Postn^MCM^*) and control (*Postn^MCM^*) mice 4 weeks after TAC or sham surgery. Data are presented as mean±SD. *P* values were calculated by two-way ANOVA with Tukey’s multiple comparison test (n=8-16 each).

**Figure S6.**
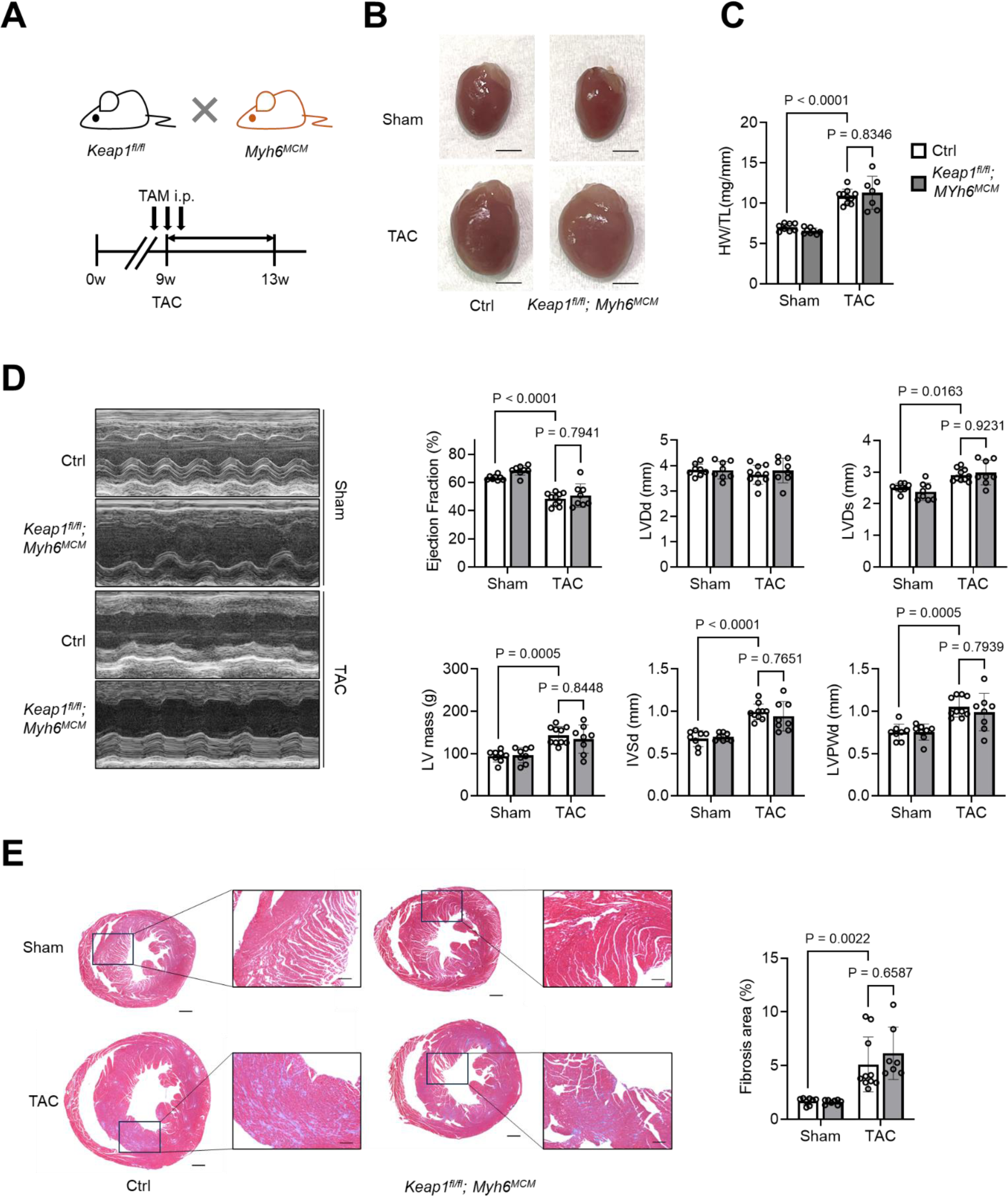
Hypertrophy and fibrosis are not affected by loss of Keap1 in cardiomyocytes. **A**, *Keap1^fl/fl^* mice were crossed with *Myh6*-*MerCreMer* (*Myh6^MCM^*) mice to generate tamoxifen-inducible cardiomyocyte-specific *Keap1* knockout mice. Adult 9-week-old male mice were subjected to transaortic constriction (TAC) or sham surgery and Cre activity was induced by intraperitoneal injection followed by feeding with tamoxifen. The mice were sacrificed 4 weeks after TAC surgery. **B**, Representative photographs of hearts from cardiomyocyte-specific *Keap1*-deficient (*Keap1^fl/fl^*, *Myh6^MCM^*) and control (*Myh6^MCM^*) mice 4 weeks after TAC or sham surgery. Scale bars: 5 mm. **C**, Quantitative analysis of heart weight to tibial length ratio. **D**, Representative echocardiographic M-mode images of left ventricle, ejection fraction (EF), left ventricular (LV) mass, and wall thickness. **E**, Representative photomicrographs of Masson’s trichrome stained transverse sections and quantification of interstitial fibrosis area of TAC-operated hearts. Scale bars: 500 µm. Data are presented as mean±SD. *P* values were calculated by two-way ANOVA with Tukey’s multiple comparison test (n=7-10 each).

